# Antidepressants interact with sex steroid receptors and their intracellular signaling components

**DOI:** 10.64898/2026.03.17.712321

**Authors:** Shokouh Arjmand, Masoud Rezaei, Donato Sardella, Claudia R. Cecchi, Rachele Rossi, Christian B. Vægter, Heidi K. Müller, Jayashree Sahana, Morten S. Nielsen, Anne M. Landau, Ulf Simonsen, Steffen Sinning, Gregers Wegener, Sâmia Joca, Caroline Biojone

**Author notes:** Correspondence to: Shokouh Arjmand, Sâmia Joca & Caroline Biojone, Palle Juul-Jensens Boulevard 99, A601/A701, Translational Neuropsychiatry Unit, Department of Clinical Medicine, Aarhus University, 8200 Aarhus N, Denmark & Høegh-Guldbergs Gade 10, Building 1115, 8000 Aarhus C, Denmark.

## Abstract

There is growing interest in understanding how hormonal signaling pathways contribute to the pathophysiology of mood disorders, based on the premise that fluctuations in sex hormones influence mood, a relationship particularly evident in conditions such as premenstrual dysphoric disorder, prenatal depression, postpartum depression, and perimenopausal depression. Estrogen receptor alpha (ERα) is predominantly localized in the nucleus, but can also be associated with the cell membrane, thus mediating a broad range of genomic and non-genomic effects through distinct intracellular pathways. By employing a combination of computational simulations and in vitro biochemical and cell-based assays, we systematically evaluated the potential binding and functional interactions of antidepressant compounds with ERα. Our results provide compelling evidence that antidepressants may not only affect classical monoaminergic targets but also modulate hormone receptor activity, particularly that of ERα. These findings are consistent with the hypothesis that ERα plays an important role in mood regulation and highlight it as a potential therapeutic target. Moreover, this work raises the possibility that the clinical efficacy of certain antidepressants may, at least in part, derive from their capacity to influence estrogen receptor-mediated signaling.

**Significance statement:** Clinical observations suggest a link between female sex hormones and mood, highlighted by the higher prevalence of depression in women and increased vulnerability to depression during hormonal fluctuations. Here, we report that structurally diverse conventional and rapid-acting antidepressants directly interact with estrogen receptor alpha (ERα). This interaction is associated with rapid intracellular signaling in cellular models. These findings indicate that, alongside their conventional targets, antidepressants may also engage sex steroid receptor components and signaling. This work broadens our basic understanding of antidepressant pharmacology at the cellular level, offering an additional perspective that may inform future research into the biological mechanisms of mood disorders and suggest a framework for developing targeted therapies for hormone-associated depressive disorders.

## Introduction

Major depressive disorder (MDD) stands out as one of the primary contributors to global disability, surpassing other significant health concerns such as ischemic heart disease, cerebrovascular disorders, diabetes, Alzheimer’s disease and other dementias, neoplastic conditions, and infectious diseases ^1^. In terms of years lived with disability, depressive disorders rank second, and they were ranked 13^th^ in disability-adjusted life-years in 2019 ^1,2^. It is believed that depression will have become the leading cause of global disease burden by 2030 ^3^.

MDD disproportionately affects women, with twice as many cases reported in women as in men ^4–6^. Moreover, women tend to experience more severe depressive symptoms, often demonstrating an earlier onset of the disease as well as more recurrent episodes with a prolonged duration that leads to an overall poorer quality of life ^7,8^. In addition, premenstrual dysphoric disorder (PMDD) manifests with depressive symptoms and mood swings, aligning with similar observations in postpartum, prenatal, and perimenopausal depression. Beyond biopsychosocial factors and the roles played by genetic and epigenetic regulators that cannot be overlooked, a contributing factor potentially explaining the sex-related differences in MDD and depressive disorders may be hormonal disparities and their associated biological consequences ^4–6,9,10^. This collective evidence, coupled with the nearly doubled incidence of depression following puberty, further alludes to a potential causal link between sex hormones, especially estrogen and progesterone, and depressive symptomatology ^11,12^.

Estrogen is *de novo* synthesized from cholesterol in the brain by neurons and astrocytes as one of the neurosteroids ^13,14^. There is a consensus that both neuron- and astrocyte-derived estrogens are crucial in promoting neuroprotection and regulating cognitive functions ^15,16^. Estrogen influences various aspects of neuronal function, including synaptic plasticity, spine synapse formation, migration, differentiation, and the survival of newborn neurons. Additionally, estrogen contributes to the proliferation of neural stem cells and the integrity of the blood-brain barrier ^13,17–23^. These regulatory effects of estrogens are achieved through their interaction with different estrogen receptors (ERs) in the brain, although some instances of ligand-independent activation of estrogen receptors have also been reported ^24^.

Interestingly, selective serotonin reuptake inhibitors (SSRIs), which typically require several weeks of continuous administration to produce a response (50 % symptoms reduction), can alleviate the depresive symptoms associated with PMDD when administered intermittently during the luteal phase ^25–27^. These findings suggest a potential interaction between antidepressants and sex steroid signaling, particularly involving estrogenic mechanisms, in the brain. The present study focuses specifically on estrogen receptor alpha (ERα), based on the hypothesis that membrane-associated estrogen signaling may play a preferential role in mediating faster antidepressant responses. Nuclear-initiated ERα signaling operates over a longer timescale, as it relies on changes in gene transcription and subsequent protein turnover. In contrast, membrane-initiated ERα signaling is thought to elicit more rapid cellular responses through the activation of various kinase signaling cascades ^28,29^.

The recognition of sex differences in depression not only provides valuable insights into the underlying causes and mechanisms of the condition but also has significant implications for refining therapeutic approaches applicable to both men and women. Given the immediate and delayed modes of action of ERα, it may emerge as a potential therapeutic target for the development of more effective antidepressants that can offer a combination of rapid onset of action and sustained antidepressant effects.

## Results

### Antidepressants activate ERα

Phosphorylation of ERα at serine 167 (Ser167) is a critical regulatory event that integrates growth-factor signaling pathways with estrogen-dependent transcription ^30,31^. Ser167, located within the AF-1 region, is targeted by several kinases, including AKT, RSK, and S6K, linking ERα to PI3K/AKT, MAPK/ERK, and mTOR signaling networks ^14^. Phosphorylation at this site enhances ERα’s transcriptional activity, stability, and chromatin binding, thereby amplifying downstream transcriptional programs associated with proliferation, survival, and metabolic adaptation ^30,31^. Ser167 phosphorylation is involved in ligand-independent receptor activation, enabling ERα to regulate gene expression ^32^. In the ligand-bound state, phosphorylation of Ser167 further modulates ERα function by enhancing synergy between AF-1 and AF-2, increasing coactivator recruitment (e.g., SRCs, p300/CBP), and extending receptor residence on chromatin ^33,34^. These effects amplify the magnitude and duration of estrogen-driven transcription and can shift promoter selectivity toward gene programs that favor cell-cycle progression and survival. Ser167 phosphorylation also reduces estrogen-induced receptor turnover, stabilizing ERα after ligand activation and lowering the effective estrogen concentration needed for transcriptional activation ^31^. Together, these ligand-dependent and ligand-independent consequences establish Ser167 as a key site for determining ERα signaling strength, estrogen sensitivity, and endocrine responsiveness.

To investigate the ability of antidepressants to modulate ERα activity, we measured phosphorylation at Ser167, using a sandwich ELISA with luminescence readout. Initially, in our experimental setup, we validated the existing findings demonstrating that estradiol induces phosphorylation of ERα at Ser167 in a concentration-dependent manner in the breast cancer cell line MCF-7, which abundantly expresses ERα but not SERT or TrkB (Cancer Cell Line Encyclopedia (CCLE), The Human Protein Atlas, and ^35^). Phosphorylation was observed within 10 minutes and 24 hours of treatment (Fig. S1A).

Various concentrations of antidepressants were tested (Fig. S1B and S1C), and we selected imipramine and S-ketamine at 10 µM to use in all subsequent experiments; this concentration falls within the clinically relevant reported brain levels of antidepressants following a treatment of adequate duration ^36–39^. Both compounds activated ERα similarly to estradiol, indicating a potential interaction of antidepressants with ERα (Fig. 1A) (in the acute treatment, the p-values were <0.0001 for estradiol and S-ketamine, and <0.01 for imipramine (Fig. 1A); in the chronic treatment, the p-values were <0.01 for estradiol and S-ketamine, and 0.0731 for imipramine (Fig. S1D)). Interestingly, the phosphorylation of ERα occurred within just 10 minutes of treatment, suggesting a rapid cellular response induced by antidepressants.

**Figure 1.**
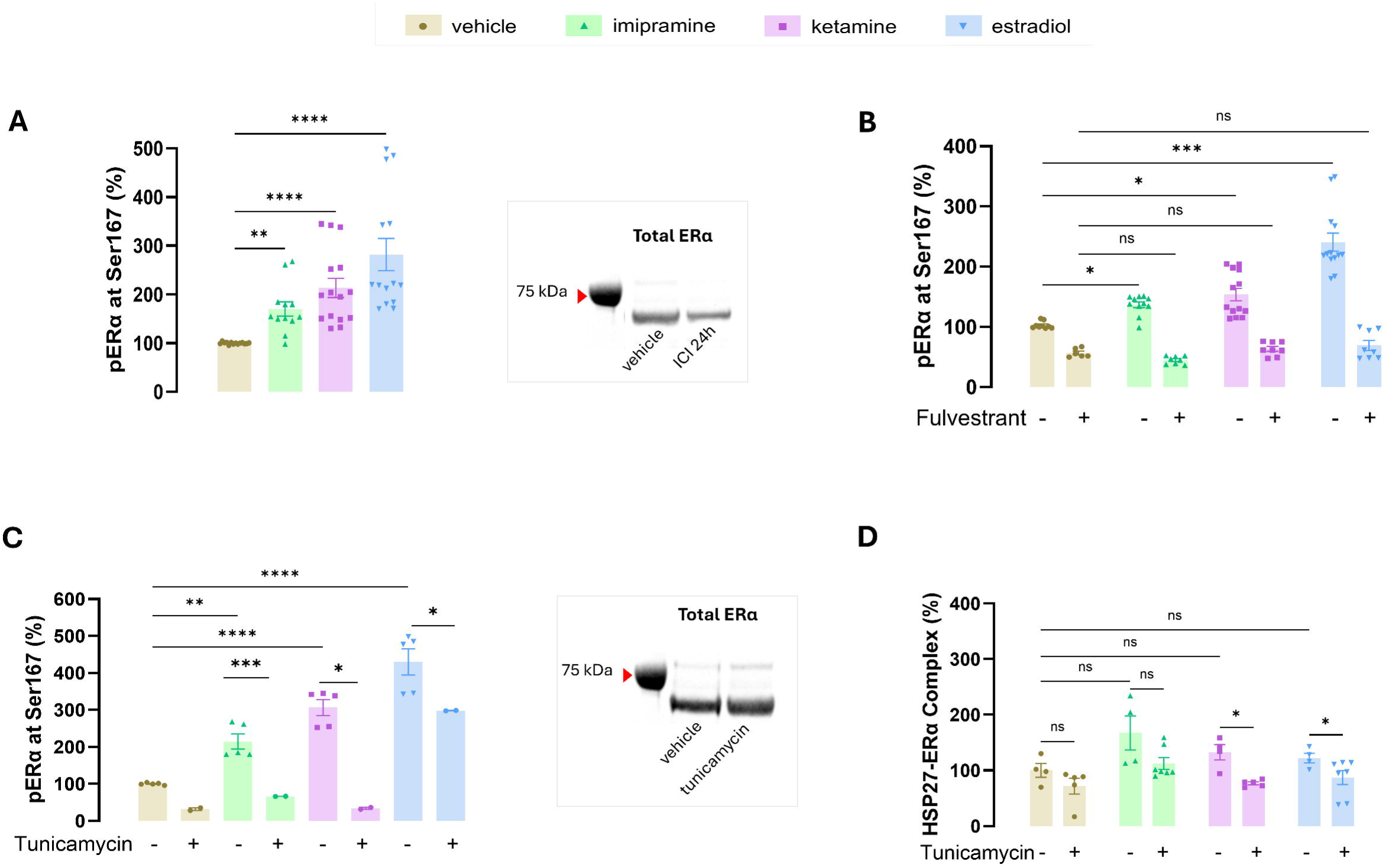
Antidepressants rapidly activate ERα. **(A)** The relative change in ERα phosphorylation at Ser167 in MCF-7 cells compared with baseline control following treatment with S-ketamine (10 µM) and imipramine (10 µM) for 10 minutes. Phosphorylation levels were measured using a sandwich ELISA with luminescence readout. Estradiol (100 nM) served as the positive control for ERα activation. (**B)** Inhibition of ERα by fulvestrant pretreatment eliminates antidepressant activation of ERα as measured by Ser167 phosphorylation. A 24-hour pretreatment of ERα with fulvestrant at 200 nM was followed by a 10-minute treatment while maintaining the concentration of ICI at 100 nM. Western blotting confirmed a lower amount of ERα protein after ICI pretreatment. (**C)** MCF-7 cells were treated with S-ketamine (10 µM), imipramine (10 µM), or estradiol (100 nM) for 10 minutes. All treatments gave rise to an increase in the level of phosphorylated Ser167 compared to the baseline. Inhibition of membrane ERα by tunicamycin (15 µM for 30 min) prior to treatment with antidepressants and estradiol reduces the effect of antidepressants and estradiol. Western blotting exhibited no change in the amount of total ERα protein after pretreatment with tunicamycin. (**D)** HSP27 in complex with ERα after pretreatment with tunicamycin (15 µM for 30 min) in various 10-minute treatment conditions. The levels of HSP27- ERα complex were measured using a sandwich ELISA with luminescence readout. Tunicamycin reduces the percentage of HSP27-ERα complex regardless of treatment conditions. The statistical significance is denoted as (ns: non-significant, *p < 0.05, **p < 0.01, *** p< 0.001, ****p < 0.0001).

As expected, pretreatment with the ERα antagonist, fulvestrant (ICI 182780), at a concentration of 200 nM for pretreatment (24h) and 100 nM during treatment (10 min), reduced basal ERα phosphorylation to approximately 50% of vehicle levels and completely abolished the rapid (10-min) increase in Ser167 phosphorylation induced by S-ketamine, imipramine, and estradiol (Fig. 1B).

A mixed-effect model revealed significant main effects of the antidepressant treatments (F (1, 68) =203.4, p<0.0001) and ICI (F (1.689, 38.27) =22.86, p<0.0001), as well as a significant interaction between the antidepressants and ICI (F (1.689, 38.27) =13.73, p<0.0001). This pattern is consistent with ERα being required either as the target for phosphorylation or as a scaffold that recruits the relevant kinases. Because fulvestrant is a selective estrogen receptor degrader (SERD) that can both degrade ERα and alter its conformation, the loss of responsiveness could reflect reduced receptor abundance (ERα depletion), conversion of residual receptor to a phosphorylation-incompetent state, or disruption of ERα-dependent scaffolding of upstream kinases.

### The rapid cellular response to antidepressants is mediated preferentially through ERα localized at the membrane

The cellular localization of ERα is integral to its biological function. The control of ERα translocation to the membrane occurs through the palmitoylation of cysteine residue 447 (Cys447) in humans ^40^. This palmitoylation process is a pivotal regulatory step influencing the functions of ERα associated with the membrane and, consequently, impacting various cellular responses mediated by ERα ^40^. To specifically assess the role of membrane ERα in the observed response to antidepressant treatment, we first inhibited palmitoylation with a brief, low concentration exposure to tunicamycin (15 µM for 30 min), thereby blocking ERα membrane localization ^41,42^.

We replicated that a 10-min treatment of MCF-7 cells with antidepressants or estradiol significantly increased ERα Ser167 phosphorylation (p<0.0001 for estradiol and S-ketamine; p<0.01 for imipramine; Fig. 1C). Pretreatment with tunicamycin attenuated the phosphorylation induced by both antidepressants and estradiol (F (7, 20) = 35.51, p<0.0001; Fig. 1C). Notably, tunicamycin reduced antidepressant-induced pERα by as much as 89% for S-ketamine (p<0.0001) and 69% for imipramine (p<0.05), whereas estradiol-induced phosphorylation was reduced by only 30% (p<0.05). This differential sensitivity to tunicamycin suggests that antidepressants primarily engage membrane-associated ERα, whereas estradiol signals through both membrane and nuclear ERα, with a predominant contribution from the tunicamycin-insensitive nuclear receptor pool. Consistent with a trafficking-specific effect, tunicamycin pretreatment did not alter total ERα protein levels, as confirmed by Western blotting (Fig. 1C).

To further examine whether the effects of the antidepressants on ERα phosphorylation were preferentially directed toward membrane-associated ERs, we investigated the interaction between ERα and heat-shock protein 27 (HSP27). HSP27 lacks the DHHC signature motif characteristic of palmitoyl acyltransferases (PATs), but it has been shown to indirectly promote ERα palmitoylation by inducing a conformational change that enhances ERα interaction with PATs ^43^. This, in turn, affects ERα trafficking to the membrane and modulates its function. Notably, HSP27 knockdown impairs membrane-associated ERα signaling without affecting its nuclear activity ^43^. We took advantage of this interaction and demonstrated that tunicamycin pretreatment reduces the overall proportion of HSP27–ERα complexes, independent of treatment conditions (main effect of tunicamycin: F (1, 32) = 17.95; p = 0.02), consistent with impaired trafficking of ERα to the membrane. No significant interaction between tunicamycin pretreatment and treatment was observed (F (1.774, 18.93) = 0.4533; p = 0.6193; Fig. 1D).

### Antidepressants alter the signaling cascade of kinases associated with membrane ERα signaling

Ligands that preferentially activate membrane-associated ERα have been shown to increase phosphorylation of mitogen-activated protein kinase (MAPK) ^44^. To determine whether antidepressants engage similar signaling pathways, we quantified phosphorylated MAPK (pMAPK) relative to total MAPK levels (Fig. 2A and 2B, respectively) following treatment with antidepressants. Both antidepressants increased the fraction of pMAPK to total MAPK (p = 0.0275 for imipramine; p = 0.0398 for S-ketamine; Fig. 2A). PI3K/Akt/MAPK is another important ERα signaling ^14^. In contrast to MAPK, no significant changes were detected in the proportion of phosphorylated Akt (pAkt) relative to total Akt across treatment conditions (F (3, 29) =1.708, p=0.1872; Fig. 2C and 2D). Notably, imipramine exhibited a trend toward reduced pAkt levels (p = 0.0826), suggesting a potential shift in downstream signaling from Akt toward MAPK activation (Fig. 2C). Pretreatment with fulvestrant abolished the antidepressant-induced increase in fraction of pMAPK (Fig. 2A). Fulvestrant pretreatment also increased overall levels of both phosphorylated and total MAPK and Akt, reflected by a significant treatment interaction (F (4.084, 29.95) =4.646, p=0.0047; Fig. 2B and 2D). In contrast, tunicamycin pretreatment selectively reduced the pMAPK response to imipramine (p= 0.0186) and showed a non-significant trend toward attenuation of S-ketamine-induced pMAPK (p= 0.0844). Representative immunoblots are shown in Fig. S2.

**Figure 2.**
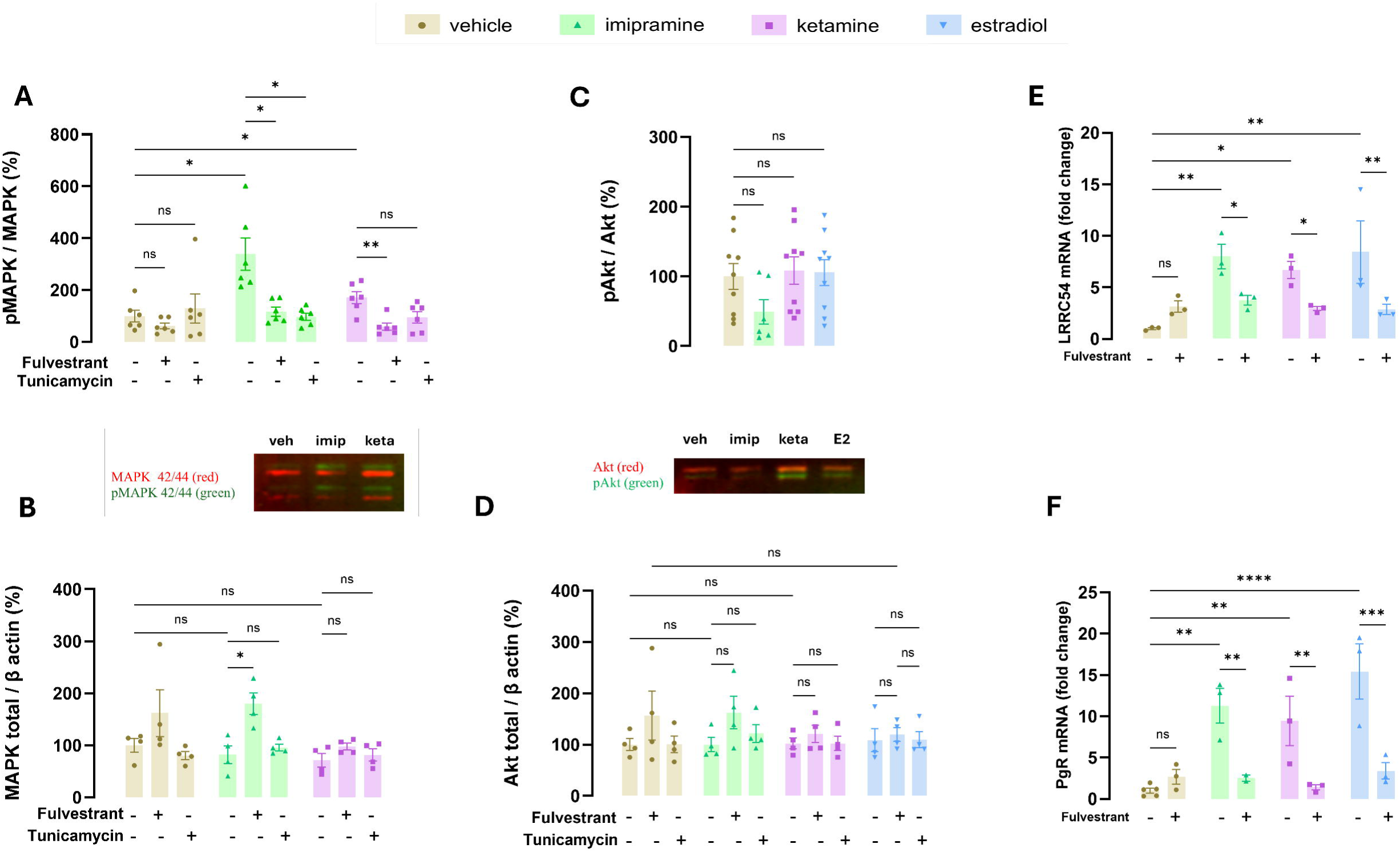
Antidepressants selectively activate MAPK signaling and induce ERα-dependent gene expression in MCF-7 cells. **(A)** Quantification of MAPK phosphorylation (pMAPK/MAPK) by Western blot densitometry following treatment with vehicle, imipramine, S-ketamine, or estradiol (E2), in the presence or absence of the ERα degrader/antagonist fulvestrant and/or tunicamycin. Antidepressant treatment significantly increased MAPK phosphorylation, indicating enhanced MAPK pathway activation. **(B)** Total MAPK levels normalized to β-actin were unchanged across treatment conditions. Representative immunoblots for MAPK and pMAPK are shown above. **(C)** Quantification of Akt phosphorylation (pAkt/Akt) revealed no significant changes following antidepressant or E2 treatment, but there was a trend for a reduction by imipramine. **(D)** Total Akt levels normalized to β-actin were unaltered. Representative immunoblots for Akt and pAkt are shown above. One imipramine-treated sample was excluded from analysis due to an outlier pMAPK/MAPK ratio exceeding 1. Data from technical and biological replicates were pooled. **(E–F)** mRNA expression of the extranuclear-initiated ERα target gene *LRRC54* **(E)** and the classical nuclear ERα target gene *PgR* **(F)**, quantified by qPCR following treatment with E2, imipramine, or S-ketamine. Antidepressants significantly increased *LRRC54* and *PgR* expression, comparable to E2 and relative to vehicle control. Pretreatment with fulvestrant attenuated these effects, indicating ERα dependency. Data are presented as mean ± SEM from three independent biological replicates (n = 3). Statistical significance is indicated as *p < 0.05, **p < 0.01, ***p < 0.001, ****p < 0.0001, and ns as not significant.

Overall, our probing of MAPK phosphorylation show that antidepressants do not significantly change the overall levels of MAPK, but that antidepressants rather shift the fraction that is phosphorylated in a manner that depends on active ERα at the cell surface.

### Antidepressants regulate genes associated with extranuclear and nuclear ERα activation

To further identify subcellular differences in how antidepressants signal via ERα we focused on the expression of a marker of extranuclear vs nuclear ERα activation. Leucine-Rich Repeat-containing 54 *(LRRC54),* also known as Tsukushi, is a gene that is reported to be preferentially induced by the activation of extranuclear ERα, whereas expression of the progesterone receptor (*PgR*) is typically associated with transcriptional activity of nuclear ERα ^44^. A 10-min treatment with S-ketamine, imipramine, or estradiol significantly enhanced the expression of *LRRC54* mRNA compared to baseline levels (p = 0.0223, 0.0049, and 0.003, respectively; Fig. 2E), consistent with rapid engagement of extranuclear ERα signaling. *PgR* expression was also significantly increased by S-ketamine, imipramine, and estradiol, relative to baseline (Fig. 2F), indicating concomitant activation of nuclear ERα-dependent transcription. Pretreatment with fulvestrant, an ERα antagonist that blocks both nuclear and extranuclear ERα signaling, abolished the antidepressant-induced upregulation of *LRRC54* (F (3, 16) =3.867, p = 0.0296), and *PgR* (F (3, 17) = 6.394, p = 0.0043). Together, these findings support the notion that both imipramine and S-ketamine may elicit rapid molecular responses through activation of extranuclear ERα signaling, while also engaging nuclear ERα-dependent transcriptional pathways.

### Antidepressants target the localization of activated ERα to the nucleus

The activation of ERα leads to receptor translocation to the nucleus. To investigate whether the antidepressants induced this hallmark of the receptor canonical signaling, MCF-7 cells were treated with vehicle, S-ketamine, imipramine, or estradiol for 10 minutes and the colocalization of phosphorylated ERα (Ser 167) (pERα) with either the nucleus or cytoplasm was explored using fluorescence microscopy. All treatment conditions significantly increased nuclear localization of pERα (p < 0.0001 for imipramine and estradiol; p = 0.0069 for ketamine, Fig 3A), consistent with the observed increase in *PgR* mRNA expression levels (Fig. 2F). In contrast, analysis of cytoplasmic pERα revealed that only imipramine significantly increased its levels (p < 0.0001), whereas no effect was observed with either ketamine or estradiol (Fig. 3B). Ligand binding to monomeric cytosolic ERα induces a conformational change that exposes the nuclear localization sequence (NLS), thereby facilitating ERα translocation into the nucleus ^45^. Since antidepressants were found to promote ERα nuclear translocation, this prompted us to ask whether they might also directly bind to ERα.

**Figure 3.**
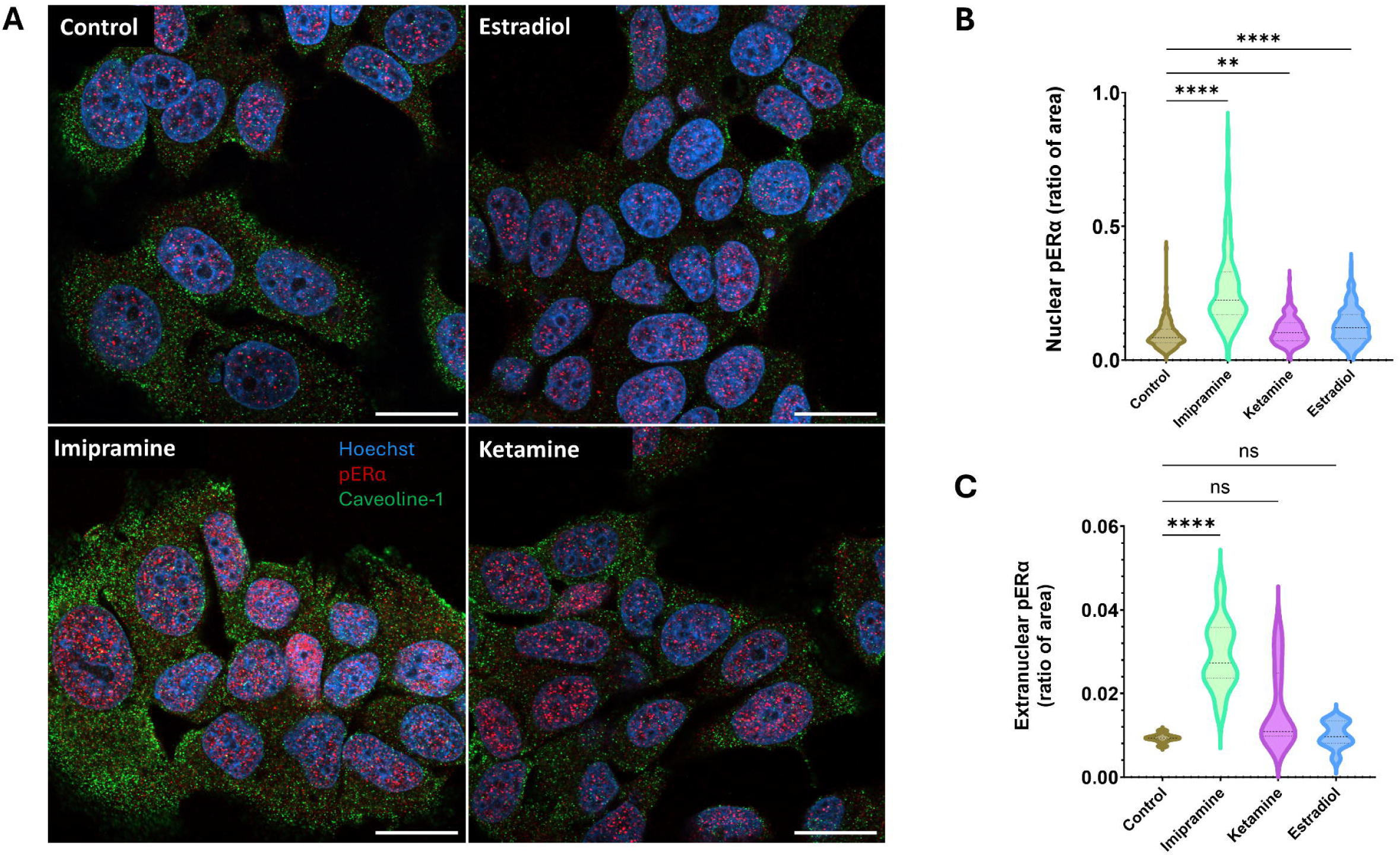
Nuclear and cytoplasmic pERα localization: statistical analysis and representative confocal images. **(A)** Ratio of pERα puncta area to nuclear area per segmented nucleus. **(B)** Ratio of pERα puncta area to cytoplasmic area per field of view. **(C)** Representative confocal fluorescence microscopy images and analyses from each experimental group, control, imipramine, ketamine, and estradiol. The red channel indicates the pERa puncta, the green channel shows Caveolin 1, and the blue channel includes Hoechst nuclear staining. Scale bars indicate 20µm. Statistical significance indicated as (ns: non-significant, *p < 0.05, **p < 0.01, ***p < 0.001, ****p < 0.0001).

### Antidepressants directly bind to estrogen receptor alpha *in silico*

To investigate the possibility of direct binding of antidepressants to ERα, molecular docking simulations were initially conducted, yielding Glide Scores (Docking Scores) for 20 selected compounds with potential or proven antidepressant activity (Table S1). The top 8 candidates were further subjected to Induced Fit Docking (IFD) simulations. The accuracy of the IFD was confirmed by superimposing the docked estradiol with the co-crystalized estradiol in the structure of 5WGD ^46^, resulting in a highly precise binding pose with a root mean square deviation (RMSD) of 0.026072 Å. The IFD analysis of estradiol revealed critical amino acids that interact with the ERα binding site, including Glu 353, Arg 394, His 524, and Phe 404 (Fig. 4A and 4B). While some antidepressants shared common interaction points with estradiol, S-ketamine and hydroxynorketamine (HNK) interacted with Gly 521 in the same binding pocket (Fig. 4C, 4D, and 5). The estimated affinities of all antidepressants predicted by SeeSAR <u>BioSolveIT</u>, ranged from picomolar to micromolar, suggesting varying degrees of binding strength (Table S2). Furthermore, the predicted topological polar surface area indicated that all tested compounds could penetrate the blood-brain barrier (BBB) and the cell membrane (Table S2).

**Figure 4.**
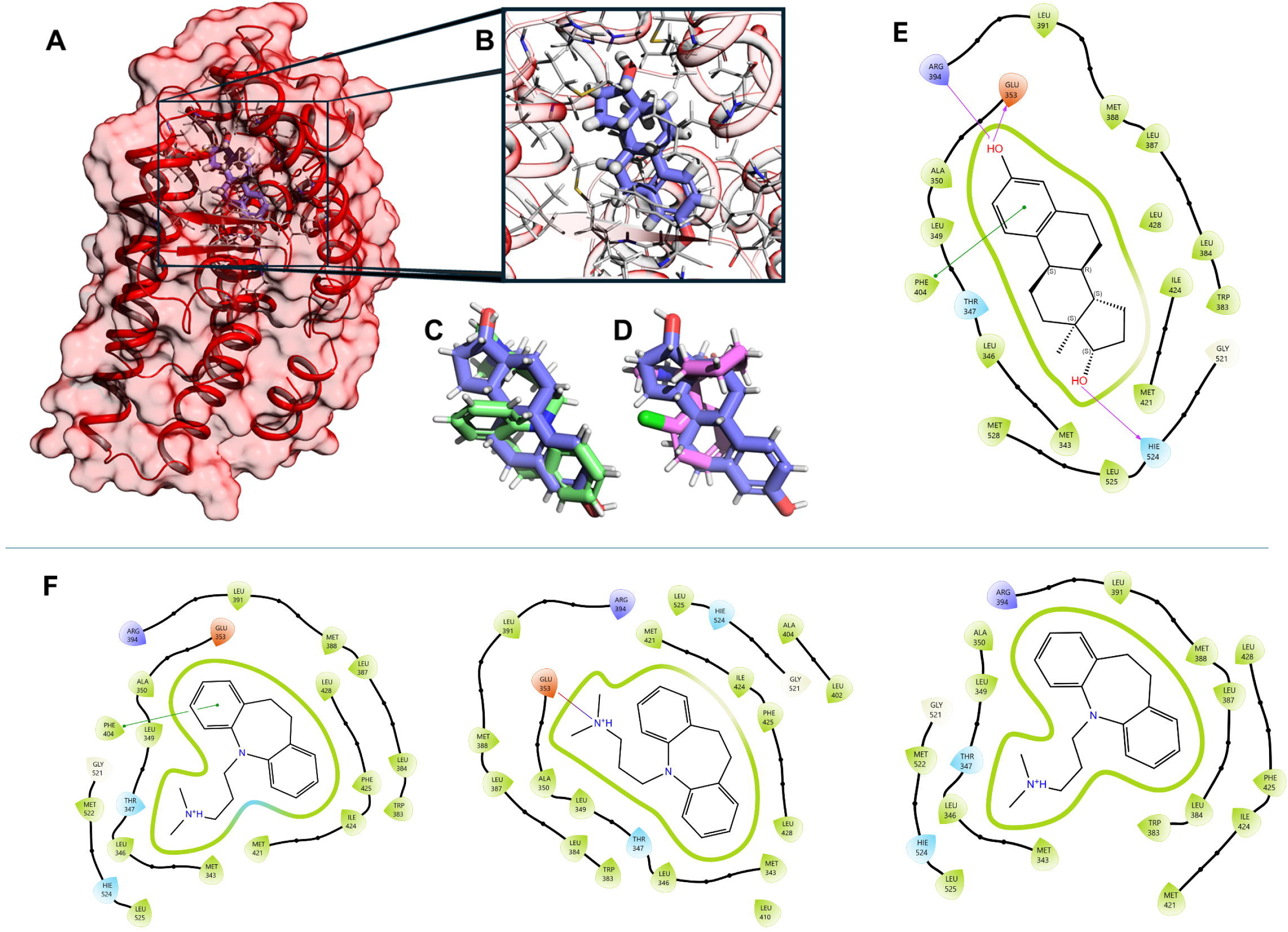
Structural basis of ligand recognition and mutational analysis in the ERα ligand-binding domain. **(A)** Overall structure of the ERα ligand-binding domain (5WGD) shown in surface (pink) and ribbon (red) representation with docked estradiol depicted as sticks in the orthosteric binding pocket. **(B)** Enlarged view of the binding site occupied by estradiol (blue) illustrating the architecture of the cavity and local protein–ligand contacts. **(C)** Structural overlay of estradiol (blue) with imipramine (green) illustrating spatial alignment within the estradiol’s primary binding pocket. **(D)** Overlay of estradiol (blue) with hydroxynorketamine (HNK) (purple) demonstrating occupancy of the same orthosteric site. **(E)** Two-dimensional interaction diagram of estradiol within the ERα binding pocket, depicting key contacts including hydrogen bonding with Glu353 and His524 and hydrophobic interactions with surrounding residues (e.g., Leu, Met, Phe) including π–π stacking interaction with Phe404. **(F)** Docking-based mutational analysis of imipramine binding. Left: wild-type receptor complex (Glide score −9.419; Emodel −47.586), showing a π–π stacking interaction between the imipramine aromatic ring and Phe404. Middle: single mutant Phe404Ala (Glide score −9.355; Emodel −47.828), in which the aromatic contact is lost but replaced by an alternative interaction with Glu353, preserving a stable binding pose. Right: double mutant Phe404Ala/Glu353Ala (Glide score −8.53; Emodel −46.058), characterized by loss of specific stabilizing contacts and a reduction in docking score, although the ligand remains accommodated within the binding pocket.

**Figure 5.**
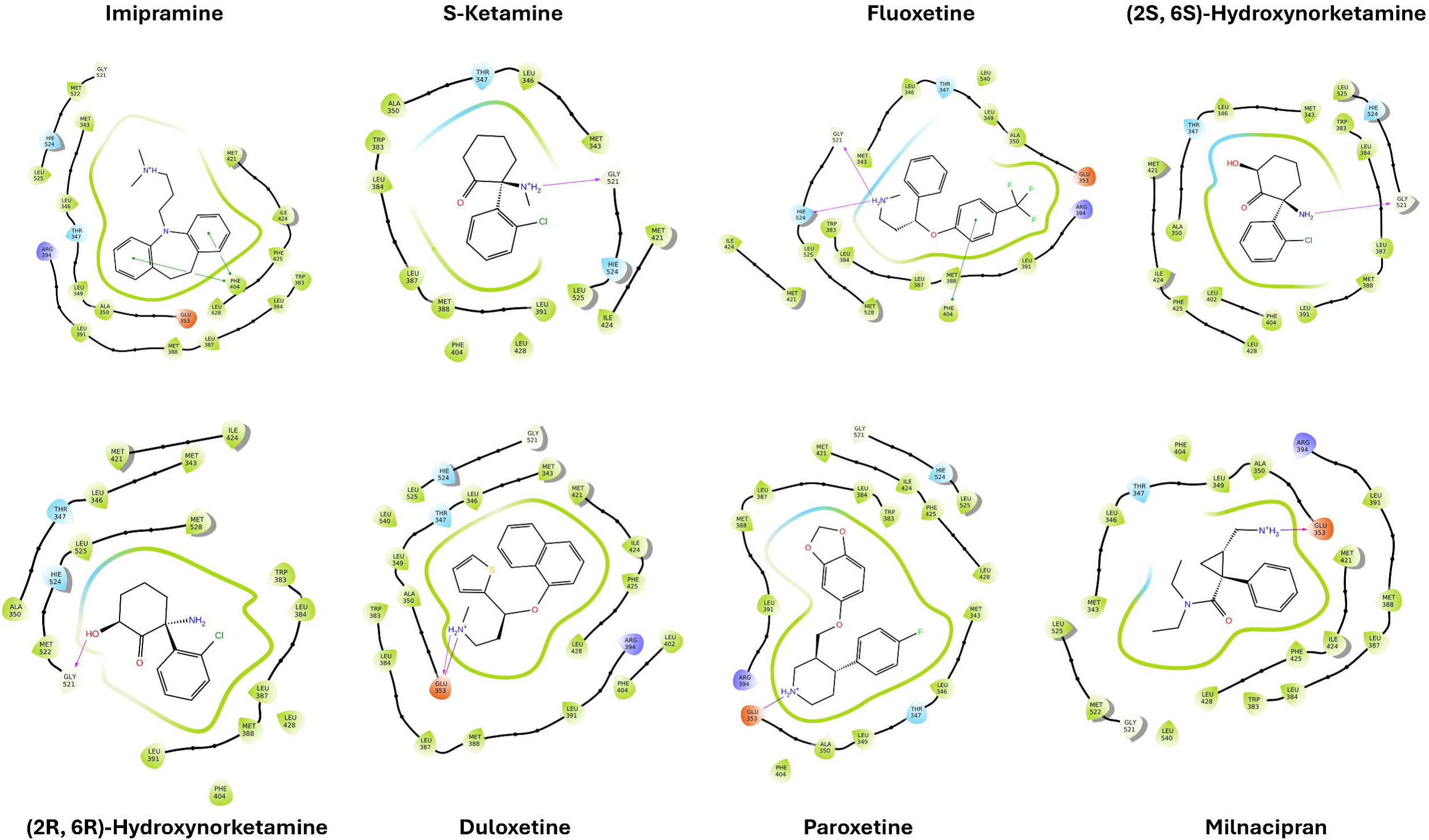
Two-dimensional representations of the interactions of various antidepressants with the ligand binding domain of ERα determined by induced-fit docking. Red circles represent negatively charged amino acids; purple circles indicate positively charged amino acids; light blue/turquoise denotes polar amino acids; and light green represents hydrophobic amino acids. The purple arrow indicates a hydrogen bond, while the double-ended dotted line represents π–π stacking interactions.

To evaluate the contribution of these amino acids in ligand binding, docking calculations were performed for imipramine as an example in the wild-type receptor and in single and double mutant models. In the wild-type complex, imipramine resulted in a Glide score of −9.419 and a Glide Emodel of −47.586, forming a π–π stacking interaction between its aromatic ring and Phe404. *In silico* mutation of Phe404 to Ala404 resulted in only a marginal change in docking score (Glide score = −9.355) and a comparable Emodel (−47.828), indicating preservation of a stable binding pose (Fig 4F). In this mutant, imipramine established an alternative interaction with Glu353, suggesting partial compensation for the loss of the aromatic contact (Fig 4F). In contrast, the double mutation of Phe404 and Glu353 led to Ala404 and Ala353 a more pronounced reduction in docking score (−8.53) and a less favorable Emodel (−46.058), accompanied by the absence of specific protein–ligand interactions (Fig 4F). Nevertheless, the Glide score remained within a favorable range, indicating that imipramine can still be accommodated within the binding pocket in the mutated receptor models. Overall, these results suggest a cooperative contribution of Phe404 and Glu353 to stabilizing the docked binding mode, while ligand association is not fully abrogated in the absence of these residues in the modeled system.

Based on these predictions, we proposed that a comparatively lower affinity for ERα might preferentially make antidepressants activate membrane ERα given their transient binding and faster dissociation from the receptor. Consequently, these antidepressants are likely to exert immediate cellular responses. To test this idea *in silico*, molecular dynamics (MD) simulations were performed to investigate the conformational dynamics and binding characteristics of four ligands, estradiol (the canonical agonist for ERα), imipramine, S-ketamine, fluoxetine, HNK, and PaPE-1 (a compound proven to preferentially bind to membrane ERα ^44^) with ERα.

The structural stability and dynamic behavior of the receptor in complex with the ligands were evaluated over 100 ns trajectories. The Ligand RMSD (Fig. 6A) revealed distinct binding stabilities; estradiol maintained the most rigid profile with fluctuations consistently below 1.5 Å, while PaPE-1 and imipramine exhibited stable trajectories equilibrating around 2.2 Å. In contrast, S-ketamine and fluoxetine showed higher mobility, with S-ketamine stabilizing at approximately 4.5 Å and fluoxetine showing a significant upward shift after 70 ns, suggesting larger conformational adjustments within the binding pocket. Despite these varying ligand dynamics, the receptor backbone RMSD (Fig. 6B) confirmed that the global structural integrity of the receptor remained intact across all systems, with values generally converging between 1.5 Å and 2.5 Å.

**Figure 6.**
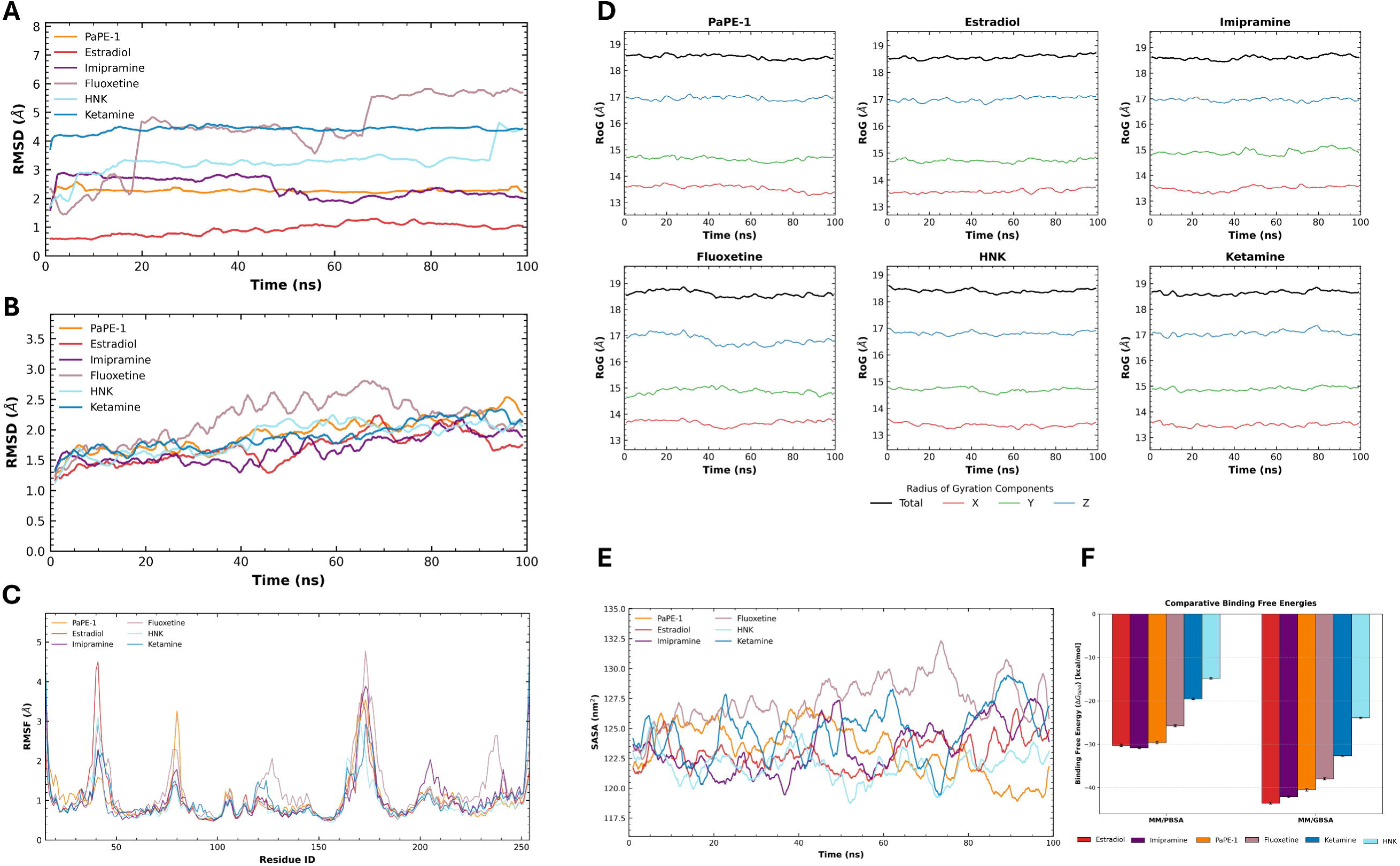
Time-dependent structural changes over 100 ns trajectories for ERα complexed with estradiol, imipramine, fluoxetine, S-ketamine, PaPE-1, and hydroxynorketamine (HNK). (**A**) Ligand root mean square deviation (RMSD) relative to the initial equilibrated structure, characterizing the stability and mobility of each compound within the binding pocket. (**B**) Protein backbone Cα RMSD, demonstrating the global structural integrity and stability of the receptor across all systems. (**C**) Cα root mean square fluctuation (RMSF) per residue, illustrating local protein flexibility and conserved dynamic patterns across the ligand-binding domain. (**D**) Radius of gyration (Rg) and its principal components (X, Y, Z), representing the maintenance of global structural compactness and shape. (**E**) Solvent accessible surface area (SASA) of the protein–ligand complexes, indicating stable exposure of the receptor-ligand surface throughout the simulation. (**F**) Visual representation of comparative binding free energies for the compounds based on MM/PBSA and MM/GBSA methods.

Local protein flexibility was assessed using the root mean square fluctuation (RMSF) of Cα atoms (Fig. 6C). The fluctuation patterns were highly conserved across all complexes, with major peaks (representing flexible loops) reaching 4–5 Å at specific residue clusters, while the core secondary structures remained stable. Global structural compactness was verified by the radius of gyration (RoG) and its principal components (Fig. 6D). The total RoG for all complexes remained stable at approximately 18.5 Å, with the X, Y, and Z components showing no significant ligand-induced anisotropic changes. The solvent accessible surface area (SASA) (Fig. 6E) fluctuated within a consistent range of 117.5 nm^2^ to 132.5 nm^2^, suggesting that the overall surface exposure and hydration of the complexes remained stable throughout the simulation period.

To estimate the binding affinities, binding free energies were calculated using both the MM/PBSA and MM/GBSA methods (Table 1 and Table 2, respectively; Fig. 6F). According to the MM/PBSA calculations, imipramine exhibited the most favorable estimated binding free energy (ΔTOTAL = - 30.79 ± 0.21 kcal/mol), followed closely by estradiol (ΔTOTAL = -30.25 ± 0.27 kcal/mol) and PaPE-1 (ΔTOTAL = -29.58 ± 0.28 kcal/mol). Fluoxetine showed a moderate affinity (ΔTOTAL = - 25.75 ± 0.26 kcal/mol), while S-ketamine and HNK showed comparatively less favorable binding energies. The dominant favorable contributions to binding for all ligands were from van der Waals interactions (ΔVDWAALS ranging from -30.36 to -45.05 kcal/mol) and electrostatic interactions (ΔE_EL_ ranging from -2.98 to -16.11 kcal/mol). The polar solvation term (ΔE_PB_) was consistently unfavorable, opposing binding, while the nonpolar solvation term (ΔE_NPOLAR_) provided a smaller, favorable contribution.

**Table 1:** MM/PBSA Binding Free Energy Components for Ligand Binding to ERα (kcal/mol)

| Ligand | $\Delta\text{VDWAALS}$ | $\Delta\text{E}_{\text{EL}}$ | $\Delta\text{E}_{\text{PB}}$ | $\Delta\text{E}_{\text{NPOLAR}}$ | $\Delta\text{G}_{\text{GAS}}$ | $\Delta\text{G}_{\text{SOLV}}$ | $\Delta\text{TOTAL}$ |
| --- | --- | --- | --- | --- | --- | --- | --- |
| Estradiol | $-43.38 \pm 0.18$ | $-11.69 \pm 0.35$ | $28.31 \pm 0.17$ | $-3.49 \pm 0.00$ | $-55.07 \pm 0.31$ | $24.82 \pm 0.17$ | $-30.25 \pm 0.27$ |
| Imipramine | $-45.05 \pm 0.16$ | $-4.87 \pm 0.12$ | $22.94 \pm 0.14$ | $-3.81 \pm 0.01$ | $-49.92 \pm 0.23$ | $19.13 \pm 0.14$ | $-30.79 \pm 0.21$ |
| Fluoxetine | $-43.65 \pm 0.20$ | $-4.62 \pm 0.22$ | $26.50 \pm 0.25$ | $-3.99 \pm 0.01$ | $-48.27 \pm 0.30$ | $22.52 \pm 0.25$ | $-25.75 \pm 0.26$ |
| S-ketamine | $-36.23 \pm 0.13$ | $-2.98 \pm 0.11$ | $22.68 \pm 0.18$ | $-2.97 \pm 0.00$ | $-39.21 \pm 0.17$ | $19.71 \pm 0.18$ | $-19.51 \pm 0.20$ |
| PaPE-1 | $-41.67 \pm 0.18$ | $-16.11 \pm 0.43$ | $31.75 \pm 0.25$ | $-3.55 \pm 0.00$ | $-57.78 \pm 0.39$ | $28.20 \pm 0.25$ | $-29.58 \pm 0.28$ |
| HNK | $-30.36 \pm 0.15$ | $-8.61 \pm 0.42$ | $27.17 \pm 0.34$ | $-3.01 \pm 0.01$ | $-38.97 \pm 0.39$ | $24.16 \pm 0.34$ | $-14.81 \pm 0.19$ |

$\Delta\text{VDWAALS}$ , van der Waals contribution from molecular mechanics; $\Delta\text{E}_{\text{EL}}$ , electrostatic energy from molecular mechanics; $\Delta\text{E}_{\text{PB}}$ , electrostatic contribution to the solvation free energy calculated by the Poisson-Boltzmann (PB) method; $\Delta\text{E}_{\text{NPOLAR}}$ , nonpolar contribution to the solvation free energy; $\Delta\text{G}_{\text{GAS}}$ , total gas-phase energy calculated as $\Delta\text{VDWAALS} + \Delta\text{E}_{\text{EL}}$ ; $\Delta\text{G}_{\text{SOLV}}$ , total solvation free energy calculated as $\Delta\text{E}_{\text{PB}} + \Delta\text{E}_{\text{NPOLAR}}$ ; $\Delta\text{TOTAL}$ , final estimated binding free energy calculated as $\Delta\text{G}_{\text{GAS}} + \Delta\text{G}_{\text{SOLV}}$ ; values represent the Average ± SEM calculated over 161 snapshots from the 100 ns production molecular dynamics trajectory; all units are reported in kcal/mol.

**Table 2:**
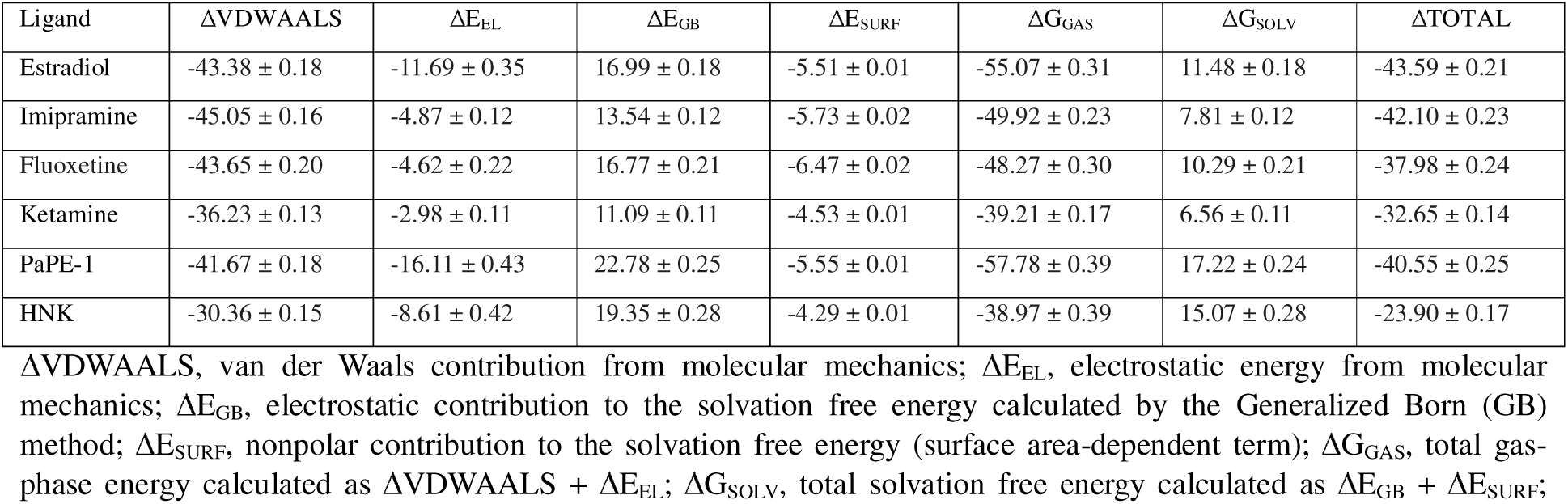
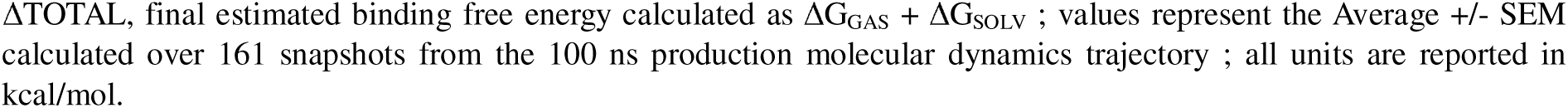
MM/GBSA Binding Free Energy Components for Ligand Binding to ERα (kcal/mol)

| Ligand | $\Delta\text{VDWAALS}$ | $\Delta\text{E}_{\text{EL}}$ | $\Delta\text{E}_{\text{GB}}$ | $\Delta\text{E}_{\text{SURF}}$ | $\Delta\text{G}_{\text{GAS}}$ | $\Delta\text{G}_{\text{SOLV}}$ | $\Delta\text{TOTAL}$ |
| --- | --- | --- | --- | --- | --- | --- | --- |
| Estradiol | $-43.38 \pm 0.18$ | $-11.69 \pm 0.35$ | $16.99 \pm 0.18$ | $-5.51 \pm 0.01$ | $-55.07 \pm 0.31$ | $11.48 \pm 0.18$ | $-43.59 \pm 0.21$ |
| Imipramine | $-45.05 \pm 0.16$ | $-4.87 \pm 0.12$ | $13.54 \pm 0.12$ | $-5.73 \pm 0.02$ | $-49.92 \pm 0.23$ | $7.81 \pm 0.12$ | $-42.10 \pm 0.23$ |
| Fluoxetine | $-43.65 \pm 0.20$ | $-4.62 \pm 0.22$ | $16.77 \pm 0.21$ | $-6.47 \pm 0.02$ | $-48.27 \pm 0.30$ | $10.29 \pm 0.21$ | $-37.98 \pm 0.24$ |
| Ketamine | $-36.23 \pm 0.13$ | $-2.98 \pm 0.11$ | $11.09 \pm 0.11$ | $-4.53 \pm 0.01$ | $-39.21 \pm 0.17$ | $6.56 \pm 0.11$ | $-32.65 \pm 0.14$ |
| PaPE-1 | $-41.67 \pm 0.18$ | $-16.11 \pm 0.43$ | $22.78 \pm 0.25$ | $-5.55 \pm 0.01$ | $-57.78 \pm 0.39$ | $17.22 \pm 0.24$ | $-40.55 \pm 0.25$ |
| HNK | $-30.36 \pm 0.15$ | $-8.61 \pm 0.42$ | $19.35 \pm 0.28$ | $-4.29 \pm 0.01$ | $-38.97 \pm 0.39$ | $15.07 \pm 0.28$ | $-23.90 \pm 0.17$ |

$\Delta\text{VDWAALS}$ , van der Waals contribution from molecular mechanics; $\Delta\text{E}_{\text{EL}}$ , electrostatic energy from molecular mechanics; $\Delta\text{E}_{\text{GB}}$ , electrostatic contribution to the solvation free energy calculated by the Generalized Born (GB) method; $\Delta\text{E}_{\text{SURF}}$ , nonpolar contribution to the solvation free energy (surface area-dependent term); $\Delta\text{G}_{\text{GAS}}$ , total gas-phase energy calculated as $\Delta\text{VDWAALS} + \Delta\text{E}_{\text{EL}}$ ; $\Delta\text{G}_{\text{SOLV}}$ , total solvation free energy calculated as $\Delta\text{E}_{\text{GB}} + \Delta\text{E}_{\text{SURF}}$ ;
ΔTOTAL, final estimated binding free energy calculated as $\Delta G_{\text{GAS}} + \Delta G_{\text{SOLV}}$ ; values represent the Average ± SEM calculated over 161 snapshots from the 100 ns production molecular dynamics trajectory ; all units are reported in kcal/mol.

Further energetic evaluation using the MM/GBSA method (Table 2) indicated that estradiol presented the most favorable binding free energy (ΔTOTAL = -43.59 ± 0.21 kcal/mol), followed by imipramine (ΔTOTAL = -42.10 ± 0.23 kcal/mol) and PaPE-1 (ΔTOTAL = -40.55 ± 0.25 kcal/mol). Fluoxetine followed with a binding energy of -37.98 ± 0.24 kcal/mol, while S-ketamine (ΔTOTAL = -32.65 ± 0.14 kcal/mol) and HNK (ΔTOTAL = -23.90 ± 0.17 kcal/mol) consistently showed the least favorable binding among the compounds tested. The binding of estradiol and imipramine was largely characterized by strong van der Waals contributions (ΔVDWAALS = -43.38 ± 0.18 and - 45.05 ± 0.16 kcal/mol, respectively). Notably, PaPE-1 distinguished itself with the most favorable electrostatic term (ΔE_EL_ = -16.11 ± 0.43 kcal/mol), though this was counteracted by a substantial unfavorable Generalized Born polar solvation penalty (ΔE_GB_ = 22.78 ± 0.25 kcal/mol).

### Antidepressants directly interact with ERα *in vitro*

To determine whether antidepressants can interact with ERα *in vitro*, we utilized fluoxetine and (2R,6R)-HNK. These compounds were selected because their secondary amine groups facilitate stable biotinylation while maintaining pharmacological activity ^47^. The hypothesized complexes of ERα and the biotinylated ligands were then subjected to an *in situ* proximity ligation assay (PLA) to detect colocalization, resulting in a positive result when the labeled targets are within 40 nm proximity. In support of a direct interaction between antidepressants and ERα, we detected a PLA signal for both biotinylated HNK and biotinylated fluoxetine not only in MCF-7 cells, which endogenously express ERα abundantly, but also in HEK293T cells transiently transfected with ERα (Fig. 7). As a negative control, a PLA signal was absent in non-transfected HEK293T cells, confirming the ERα-dependence of the observed proximity interactions (Fig. 7). The emergence of a PLA signal suggests that biotinylated fluoxetine and HNK are in close spatial proximity to ERα, implying potential direct or receptor-mediated binding interactions. Interestingly, most PLA puncta were localized to extranuclear and perinuclear regions. This spatial distribution points to the possible involvement of extranuclear, likely monomeric ERα, which has been implicated in rapid, non-genomic signaling pathways.

**Figure 7.**
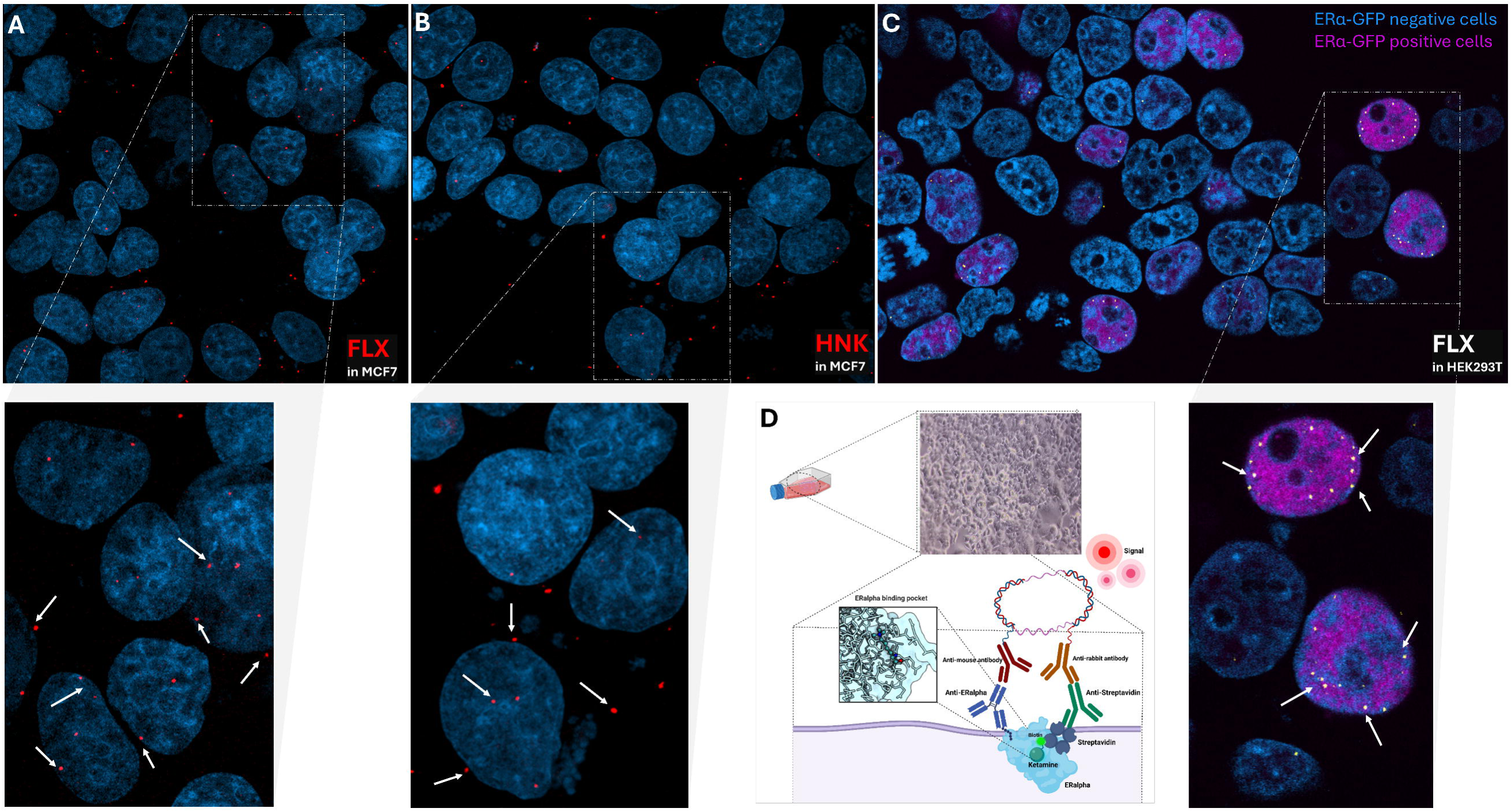
In situ PLA demonstrates close proximity between biotinylated fluoxetine as well as HNK and ERα (red dots) in MCF-7 cells and transfected HEK293T cells (magenta), while no PLA signal was detected in HEK293T cells not expressing ERα. ERα-transfected cells are tagged with GFP and are represented in magenta. Blue represents DAPI staining. Arrows point to the PLA signal.

### Antidepressants activate ERα in primary neurons

To determine whether the rapid ERα signaling observed in MCF-7 cells also occurs in neuronal tissue, we conducted complementary experiments using cultured rodent primary neurons to clarify the physiological relevance of the pathway and its potential implications for antidepressant action within the central nervous system. The experiment aimed to evaluate the responsiveness of neuronal ERα to rapid modulation by antidepressants and estradiol, to provide insight into whether similar mechanisms and machinery are active in the brain.

Imipramine, ketamine, and estradiol significantly increased pERα (p = 0.0393, 0.0005, and <0.0001, respectively), and pretreatment with tunicamycin markedly attenuated this effect (Fig. 8), implying that the same phenomenon also occurs in brain cells.

**Figure 8.**
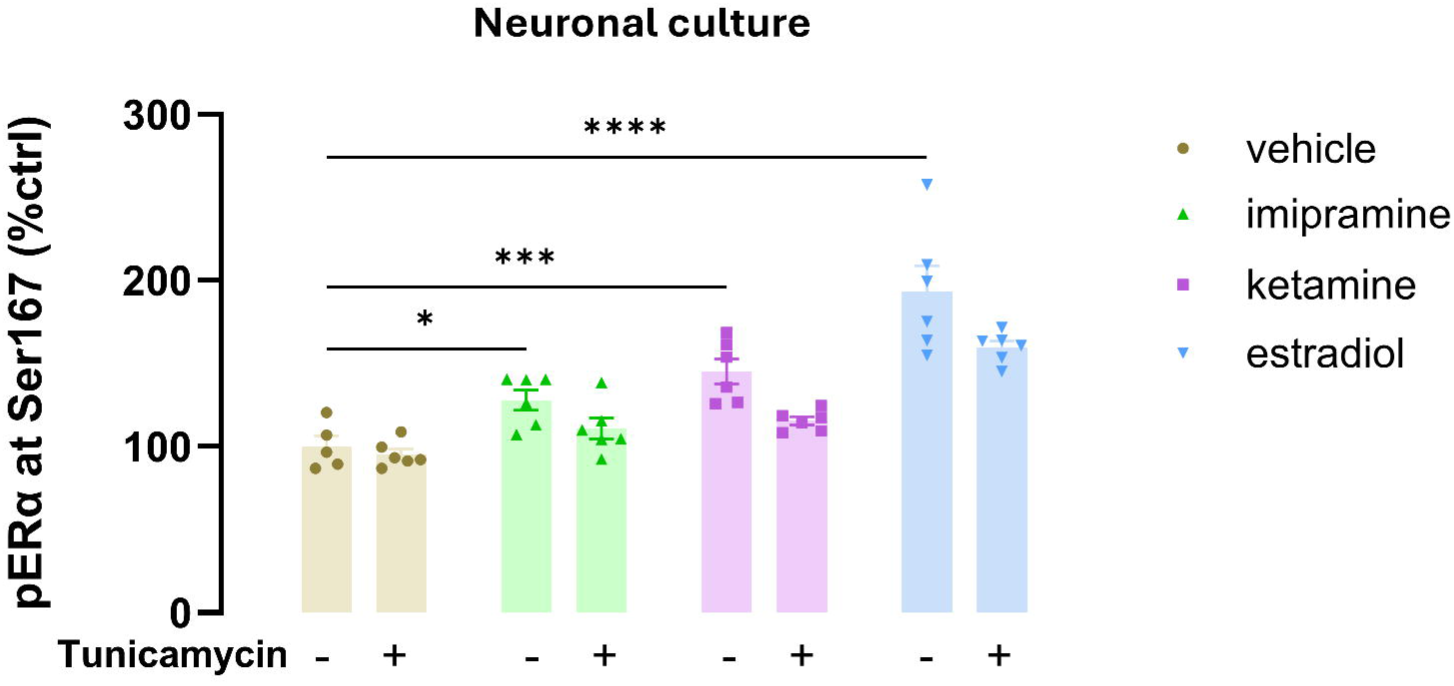
Cultured rodent primary neurons were treated with imipramine (10 μM), ketamine (10 μM), or estradiol (100 nM) for 10 minutes. All treatments induced a significant increase in pERα compared with the vehicle-treated control. This rapid phosphorylation response mirrors that observed in MCF-7 cells and demonstrates that neuronal ERα is responsive to both antidepressants and estradiol. One vehicle data point was excluded as an outlier based on an outlier test. Data represent mean ± SEM; Statistical significance indicated as (ns: non-significant, *p < 0.05, **p < 0.01, ***p < 0.001, ****p < 0.0001).

## Discussion

The current study was motivated by the clinical observation that hormonal fluctuations are closely linked to mood disturbances, as seen in disorders such as premenstrual dysphoric disorder, perinatal depression, and perimenopausal depression. These conditions suggest a biological interplay between sex steroid signaling and mood regulation. Building on this premise, we investigated whether antidepressants may exert part of their effects through interaction with ERα, a key mediator of estrogenic signaling. By integrating computational and molecular assays, our findings provide initial evidence that antidepressants can engage with ERα and modulate its subcellular localization, offering a potential mechanistic link between antidepressant action and hormone receptor pathways.

Our findings reveal an increased phosphorylation of ERα at Ser167 following antidepressants treatment, mirroring a similar response induced by estradiol in MCF-7 cells. This aligns with the established literature, showing that phosphorylation of ERα is enhanced upon estradiol binding ^48,49^. Phosphorylation of serine residues of ERα is important as it contributes to the recruitment of coactivators and activation of the receptor. Among the various phosphorylation sites on ERα, Ser167 has been identified as one of the major sites phosphorylated upon estradiol binding ^33^. Ser118, Ser104, and Ser106 are other important phosphorylation sites responsive to estradiol binding ^34^. Despite ongoing controversies regarding the phosphorylation of Ser167 in response to estradiol binding ^49,50^, our study provides clear evidence of dose-dependent phosphorylation of Ser167 following treatment with estradiol. Additionally, evidence suggests that Ser167 can be phosphorylated by various kinases, including Akt, RSK, and, especially, MAPK, further adding complexity to the regulatory mechanisms ^51–53^.

Detecting endogenous membrane ERα using standard immunofluorescence approaches is typically challenging, unlike exogenously expressed extranuclear ERα, which can be detected using antibodies against the DNA-binding domain, hinge, or carboxy-terminal ^54,55^. A study focusing on the COOH terminus of ERα successfully identified over 90% of membrane ERα, suggesting that membrane and nuclear ERα possess distinct conformations and antigen accessibility ^56^. We utilized SRA-1010, a monoclonal antibody targeting the ERα C-terminus, to specifically capture membrane ERα. While our findings show consistent receptor activation across both 10-minute and 24-hour treatments, these results could be subjected to antibody specificity. Because SRA-1010 predominantly detects extranuclear and perinuclear ERα (with only 4 ± 2% nuclear detection observed in a 30-minute treatment), we likely only captured the extranuclear fraction of the receptor

The rapid activation of ERα by antidepressants (10 minutes), together with our observation that antidepressants affect the expression of genes linked to extranuclear ERα signaling and enhance the activity of kinases associated with membrane ERα activation, implies the involvement of membrane ERα. This interpretation is further supported by the diminished cellular effects of antidepressants, including the positive control estradiol, after pretreatment with tunicamycin. Importantly, tunicamycin did not alter the total amount of ERα, indicating a selective disruption of ERα palmitoylation, likely mediated in part by reduced HSP27 binding to ERα. This observation is of importance, as membrane-initiated signaling plays a crucial role in peripheral cellular responses, particularly in endothelial cells and specific neural regions where genomic signaling is less prominent. It has been established that in regions with robust genomic signaling, membrane ERα has been reported to contribute less substantially to downstream responses *per se* ^55,57–59^. While most of our experiments and assays were performed in MCF-7 cells, in which genomic signaling is robust, potential variations in ERα signaling across different tissues should be considered. This is particularly relevant in the context of antidepressant action in the brain, where membrane-initiated signaling may play a more dominant role. This highlights the need for caution when interpreting these findings, as cellular responses may differ in brain tissues with distinct genomic signaling characteristics. Nevertheless, to confirm the relevance of our observations in a neural context, we demonstrated that antidepressants similarly induce ERα phosphorylation in cultured primary neurons, an effect that was likewise abolished by tunicamycin pretreatment, as previously detected in MCF-7 cells.

To assess whether the rapid response induced by antidepressants is abolished by ERα inhibition, we used fulvestrant, which is a cell-permeable estradiol derivative capable of functioning as an estrogen antagonist. Fulvestrant reduced ERα phosphorylation to around 50% of basal levels across all conditions and abolished the increased phosphorylation induced by antidepressants and estradiol, indicating that ERα is required for these compounds’ observed mode of action on the receptor. Fulvestrant, classified as a SERD, impedes receptor dimerization by binding to ERα monomers. This binding leads to the inactivation of both AF1 and AF2 ^60^. Consequently, receptor’s translocation to the nucleus is diminished, leading to ERα degradation and downregulation while leaving estrogen receptor-mediated progesterone receptor expression unaffected. This process occurs as fulvestrant forms an unstable complex with ERα ^60^. Thus, it is important to consider that the reduced phosphorylation of ERα might result from limited ERα availability for phosphorylation rather than from a lack of interaction between antidepressants, estradiol, and ERα due to inhibition.

We demonstrated that brief exposure to antidepressants, such as S-ketamine, imipramine, and fluoxetine, enhances the nuclear localization of phosphorylated ERα in MCF-7 cells, a pattern resembling that of estradiol. This observation is in line with the increased *PgR* expression after treatment with antidepressants. Interestingly, imipramine also increased cytoplasmic levels of phosphorylated ERα, suggesting it may differentially influence both genomic and non-genomic ERα signaling. In addition, proximity ligation assays using biotinylated fluoxetine and HNK revealed interactions with ERα in both native ERα-expressing MCF-7 cells and HEK293T cells transfected with ERα. The absence of PLA signal in non-transfected HEK293T cells confirms the specificity of these interactions to ERα, with signal localization predominantly in extranuclear and perinuclear regions, aligning with the hypothesis that antidepressants may engage extranuclear monomeric ERα involved in rapid, non-genomic signaling cascades. These non-traditional pathways have been increasingly recognized in neuropsychiatric contexts and may represent a previously underappreciated mechanism of antidepressant action.

These findings suggest that certain antidepressants may modulate ERα signaling, potentially acting as ligands or allosteric modulators that influence its activation and intracellular trafficking. The observed nuclear translocation of phosphorylated ERα in response to antidepressants parallels mechanisms typically associated with endogenous estrogen, indicating that these drugs may partially engage hormone-related signaling pathways. These findings not only strengthen the mechanistic link between antidepressant action and estrogen receptor signaling but also raise the possibility that part of the therapeutic effects of antidepressants may arise from their engagement with sex hormone receptor signaling pathways, offering a novel perspective particularly relevant for hormonally sensitive mood disorders such as postpartum or perimenopausal depression. Importantly, the direct engagement of antidepressants with ERα does not inherently restrict their therapeutic efficacy to females. Estrogen receptors are widely distributed in both male and female brains ^14,61,62^, and act as critical regulators of intracellular signaling and neuroplasticity. By directly modulating ERα, antidepressants can effectively trigger the necessary signaling cascades for antidepressant action in both sexes. While sex-differences in response to certain antidepressants exist ^63,64^, we have previously demonstrated in a preclinical model that the rapid antidepressant effects of S-ketamine are sustained regardless of sex or estrous cycle phase ^65^.

Furthermore, our computational analyses provided molecular-level insight into the potential for direct interaction between antidepressants and ERα. Molecular docking and induced fit simulations revealed that several antidepressants, including ketamine, fluoxetine, duloxetine, citalopram, bupropion, psilocin, and imipramine, can occupy the ERα ligand-binding pocket, with some sharing interaction residues with estradiol. Notably, antidepressants displayed binding free energies comparable to estradiol and PaPE-1 in both MM/PBSA and MM/GBSA calculations. These findings suggest that antidepressants may exhibit sufficient binding affinity and receptor engagement to modulate ERα activity, possibly mimicking or partially overlapping with endogenous estrogen signaling.

Estradiol exemplifies canonical ERα activation, stabilizing a compact, low-flexibility receptor conformation with favorable van der Waals and electrostatic interactions. This energetic and structural stabilization is consistent with its ability to promote both genomic transcription and non-genomic signaling. This combination of optimized packing, limited solvent exposure, and structural rigidification is consistent with stabilization of the AF-2 surface and helix 12 positioning required for coactivator recruitment and genomic transcriptional signaling.

PaPE-1, in contrast, binds ERα with affinity comparable to estradiol but induces increased conformational flexibility and solvent exposure. Its strong gas-phase electrostatic interactions are counterbalanced by polar solvation penalties, yielding a dynamically permissive receptor state. This energetic–dynamic signature reflects a ligand that binds strongly yet promotes a more conformationally permissive ERα ensemble. Such a state may be expected to reduce sustained AF-2 stabilization while maintaining receptor engagement in signaling complexes. Our simulation findings provide structural support for the membrane-preferential, non-genomic signaling profile attributed to PaPE-1 and illustrate how strong intrinsic binding can coexist with increased conformational entropy to bias signaling outcomes.

Imipramine and fluoxetine partially recapitulate this behavior. Both ligands exhibited total binding free energies comparable to estradiol and PaPE-1, largely driven by favorable van der Waals interactions rather than strong electrostatic anchoring.

Molecular dynamics revealed moderate ligand mobility and increased receptor flexibility relative to estradiol, along with modest increases in solvent exposure. These features indicate that although imipramine and fluoxetine can stably occupy the ligand-binding pocket, they do not enforce the same degree of conformational constraint required for canonical genomic signaling. Instead, their interaction profile suggests stabilization of a dynamically flexible receptor state. Such a state may be permissive for non-genomic signaling or context-dependent modulation but may lack the structural features typically associated with robust transcriptional activation.

In contrast, S-ketamine and HNK demonstrated substantially weaker binding free energies, elevated ligand RMSD, increased receptor flexibility, and expanded solvent exposure, suggesting transient or opportunistic engagement with the ERα ligand-binding domain or multiple binding poses.

Furthermore, the physicochemical properties of all compounds supported their ability to penetrate the BBB and cell membranes, aligning with their known central nervous system activity and raising the possibility of ERα modulation *in vivo*. The observation that antidepressants shared dynamic and energetic profiles with estradiol and PaPE-1, despite their distinct pharmacological classifications, raises questions about the broader structural criteria for ERα activation beyond classical steroid scaffolds. Given PaPE-1’s established preference for membrane ERα ^44^, and the comparatively lower binding energy ^44^, it is plausible that transient or lower-affinity binding may preferentially engage membrane-localized ERα, promoting rapid, non-genomic signaling.

The ligand-binding domain of ERα is a complex allosteric signaling domain that commonly possesses 11 α-helices and four β-strands that fold into three parallel layers to form an alpha-helical sandwich. Such folding creates a hydrophobic core (the ligand-binding pocket) at the base of the receptor that enables the accommodation of the ligand. The ligand-binding pocket is primarily composed of approximately 75% hydrophobic residues, a structural feature that confers substantial conformational plasticity and broad ligand accommodation capacity. However, it also includes essential polar residues that form crucial hydrogen bonding interactions with the ligand. These hydrogen bonds play a vital role in positioning the ligand correctly within the pocket ^66–68^. We postulated that this hydrophobicity likely facilitates the accommodability and binding of structurally diverse compounds, including non-steroidal molecules such as many antidepressants tested by enabling favorable van der Waals interactions and allowing adaptive fit within the binding cavity. Such physicochemical versatility may underlie the observed ability of chemically heterogeneous antidepressants to stably associate with ERα.

Consistent with the predominantly hydrophobic character of the ERα ligand-binding pocket, the docking results indicate that imipramine can be stably accommodated through nonpolar interactions, supplemented by specific residue contributions. In the wild-type model, π–π stacking with Phe404 appears to provide localized aromatic stabilization. However, the limited change in Glide score following a single mutation of this residue suggests that the overall hydrophobic environment of the pocket can buffer the loss of an individual aromatic contact. The emergence of an alternative interaction with Glu353 in the single mutant further supports the notion of adaptive binding within a conformationally permissive cavity that can accommodate structurally diverse ligands. In contrast, the double mutation produced a more noticeable reduction in docking score and loss of defined residue interactions, although the ligand remained favorably docked. Together, these observations align with the structural plasticity of the ERα binding domain, where distributed hydrophobic contacts and compensatory interactions may collectively sustain ligand accommodation rather than reliance on a single dominant residue. It should be noted that these interpretations are based on static docking models and do not account for receptor dynamics or solvent-mediated effects.

Although the present study establishes a novel interaction between antidepressants and ERα, the broader landscape of sex steroid signaling in mood regulation is highly complex and warrants further investigation. Progesterone and its associated receptors play a fundamental role in neuroplasticity and are the other players that can be involved in the pathophysiology of hormone-sensitive conditions, such as postpartum and perimenopausal depression. Given the structural homologies among steroid hormone receptors and the known rapid-antidepressant effects of neuroactive progesterone metabolites ^69^, it is highly plausible that PRs are also involved in these complex pharmacological networks. Future studies could investigate whether conventional and rapid-acting antidepressants similarly interact with PRs and their intracellular signaling cascades to fully map the sex-specific targets of these medications.

In summary, our research unveils mechanistic insight into the potential engagement of antidepressants with ERα signaling pathways, not merely through indirect modulation but also potentially through direct or allosteric binding. Our findings reveal that these interactions are predominantly mediated via extranuclear ERα, suggesting involvement of non-genomic ERα signaling. Given the known role of extranuclear ERα in mediating rapid cellular responses, these findings highlight a possible pathway through which antidepressants may influence intracellular signaling, independent of classical monoaminergic mechanisms.

## Materials and methods

### Compounds

Imipramine hydrochloride (Sigma-Aldrich, I7379), fluoxetine hydrochloride (LKT Labs, F4780), and S-ketamine (Pfizer, 2750 Ballerup, Denmark) were used as antidepressants and 17-β estradiol (Sigma-Aldrich, E2758) served as the positive control in the study. Tunicamycin (Thermo Scientific Chemicals, J67401.XF) and fulvestrant (ICI-182780) (Sigma-Aldrich, I4409) were used to inhibit palmitoylation and ERα, respectively.

### Cell lines and cell cultures

MCF-7 cells, MDA-MB-231 cells, HEK293T, and HEK293 cells were used across *in vitro* studies. MCF-7 breast cancer cells are of human origin and are responsive to estradiol ^70^. They also ubiquitously express ERα and progesterone receptors, and it has already been established that they contain membrane ERα ^70–74^, making them an ideal cell line for evaluating the interaction of antidepressants with both membrane and nuclear ERα. MDA-MB-231 cells were used as a negative control due to their lack of ERα expression ^72^. HEK239 cells were chosen to transiently overexpress ERα post-transfection, with non-transfected cells as their controls. Experiments were conducted using Dulbecco’s modified Eagle’s medium (DMEM), high glucose DMEM, and RPMI-1640 media, all containing 10% fetal bovine serum (FBS), 2 mM glutamine, and 1% penicillin/streptomycin. Notably, as phenol red acts as a potential agonist for ERα ^75^, experiments were repeated using phenol red-free DMEM and RPMI-1640 media as well.

For experiments using primary neuronal cultures, C57Bl6 mouse cortical brain tissue was dissected at embryonic day 16-17, following protocol by Sahu et al., 2019 ^76^, with minor modifications, and seeded in 1.9 cm^2^ wells (250,000/well), in poly-L-lysine coated plates. Neurons were kept for in Neurobasal medium supplemented with B27, L-glutamine and penicillin/streptomycin, and at 7 DIV they were treated with E2 or antidepressants added directly to the medium. Whole-cell lysates from immortalized cell lines or primary neurons were collected in NP lysis buffer (20 mM Tris-HCl, 150 mM NaCl, 50 mM NaF, 1% Nonidet-40, 10% glycerol) supplemented with 2 mM Na_3_VO_4_ and Halt protease inhibitor cocktail (Thermo Fisher Scientific).

### Western blotting

Cells were treated with compounds of interest, and whole-cell lysates were collected in NP lysis buffer as described above. A total of 15–20 μg of protein was loaded per lane after being mixed with sample buffer. The buffer consisted of NuPAGE™ LDS Sample Buffer and dithiothreitol (DTT) at a 9:1 ratio and was denatured by heating at 65°C for 15 minutes prior to loading onto NuPAGE™ Bis-Tris 4–12% Midi Protein Gels or NuPAGE™ 10% Bis-Tris Gels (Invitrogen). Precision Plus Protein™ All Blue Prestained Protein Standards (Bio-Rad) or PageRuler™ Prestained Protein Ladder (Invitrogen) were used as molecular weight markers. Electrophoresis was performed using NuPAGE™ MOPS SDS Running Buffer (Invitrogen) in a standard electrophoresis chamber. Proteins were transferred onto 0.2 μm nitrocellulose membranes using either the Trans-Blot Turbo Transfer System (Bio-Rad) or the iBlot® 2 Dry Blotting System (Invitrogen). After a 5-minute incubation in TBS, membranes were blocked for 1 hour using either Intercept™ Blocking Buffer (LI-COR Biosciences) or 5% bovine serum albumin (BSA) in TBS. Membranes were incubated overnight at 4°C with primary antibodies diluted in a 1:1 mixture of Intercept™ Blocking Buffer and TBST. The following primary antibodies were used: rabbit anti-phospho-Akt (Ser473) (Cell Signaling #4060), mouse anti-Akt (Cell Signaling #2920), rabbit anti-phospho-p44/42 MAPK (Thr202/Tyr204) (Cell Signaling #4370), mouse anti- p44/42 MAPK (Cell Signaling #9107). After washing, membranes were incubated for 1 hour at room temperature with the appropriate IRDye®-conjugated secondary antibodies (800CW goat/donkey anti-rabbit IgG or IRDye 680RD goat/donkey anti-mouse IgG, LI-COR Biosciences). Signal detection was performed using the Odyssey® CLx infrared imaging system, and band intensities were quantified using Image Studio version 6.0. Membranes were then stripped for 10 minutes and reprobed with antibodies against ERα (mouse anti-ERα (1D5)) and β-actin (mouse or rabbit Licor 926–42212/Licor 926-42210).

### Sandwich ELISA

Sandwich ELISA was used in at least three independent assays to measure phosphorylated ERα, and the interaction of ERα with HSP27. Briefly, white high binding 96-well plate (Revvity/Perkin Elmer) were initially coated with anti-ERα antibody of mouse origin (SRA1010, Enzo, ADI-SRA-1010-F), capable of capturing membrane ERα in addition to the cytosolic and a small fraction of nuclear ERα, diluted 500 times in carbonate buffer (25 mM sodium bicarbonate, 25 mM sodium carbonate, pH 9.8) overnight at 4°C. After blocking for 2 h at room temperature (RT) in 2% BSA in PBST (PBS + 0.1% Tween 20), lysates from MCF-7 cells or BSA (Sigma-Aldrich, A9647) or MDA-MB-231 cells were added and incubated overnight at 4°C with agitation. Subsequently, the plates were washed in PBST and primary antibodies, anti-pERα Ser167 (Cell Signaling (D5W3Z), #64508) (1:1000), or anti-HSP27 (Invitrogen, PA5-78010) (1:1000), all derived from rabbit and diluted in 2% BSA, were added to the plate and incubated at 4°C overnight with gentle shaking. Lastly, plates were washed three times with PBST buffer, and then goat anti-rabbit HRP-conjugated tertiary antibody (BioRad, 1705046) (1:5000) was added for 1h at RT. After adding ECL substrate (ThermoFisher Scientific, 32209), luminescence was measured (1 s/well) using a plate reader (Varioskan Lux, Thermo Scientific). Nonspecific signal (average luminescence from 8–10 blank wells) was subtracted from each sample, and data were expressed as a percentage of the vehicle-treated control.

### qPCR

MCF-7 cells were pre-cultured in phenol red-free RPMI 1640 medium supplemented with L-glutamine and 10% charcoal-stripped FBS to eliminate exogenous hormones. Cells were maintained in these hormone-depleted conditions throughout the entire experiment to ensure estrogen-receptor specificity of subsequent treatments. Cells were seeded at a density of 250,000 cells per well in 6-well plates and allowed to attach and proliferate until reaching 70–80% confluence prior to treatment. At 70–80% confluence, cells were treated for 4 hours with one of the following conditions (estradiol 10 nM, ketamine 10 µM, imipramine 10 µM, and vehicle). In a parallel experiment, MCF-7 cells were pretreated with fulvestrant (1 µM final concentration) for 2 hours to antagonize estrogen receptor activity. Fulvestrant (ICI) was added at 1 µL per well, and cells were incubated at 37°C. After 2 hours of fulvestrant pretreatment, a second treatment was administered for an additional 4 hours ^77^, using the same conditions as above. At the end of the 4-hour treatment, cells were immediately placed on ice and processed for total RNA isolation using the PARIS Kit (Thermo Fisher Scientific), according to the manufacturer’s instructions. All reagents and consumables were kept RNase-free. Quantitative real-time PCR was performed using standard protocols. The reference gene used for normalization was 36B4 (RPLP0) due to its stable expression under hormone-depleted and drug-treated conditions.

The sequences of primers for the genes studied were *LRRC54* forward, 5’-GGGCTACACGACGTTGGCT-3’; *LRRC54* reverse, 5’-GAGGTCAAGCGACTCCAGGTA-3’ ^77^; *PgR* forward, 5’-AGGTCTACCCGCCCTATCTC-3’; *PgR* reverse, 5’-TCCCACAGGTAAGGACACCA-3’ ^78^; *36B4* forward, 5’-CGACCTGGAAGTCCAACTAC-3’; *36B4* reverse, 5’-ATCTGCTGCATCTGCTTG-3’; and were all provided by Eurofins.

### Immunofluorescence staining for colocalization assay

MCF-7 cells were cultured on glass coverslips and processed for immunofluorescence to assess the colocalization of pERα, caveolin-1, and nuclei (Hoechst). After fixation in 4% paraformaldehyde (PFA), coverslips were rinsed with Dulbecco’s phosphate-buffered saline (DPBS). Blocking was performed for 1 hour at RT using 5% BSA in DPBS supplemented with 0.05% Tween®-20 and 0.1% Triton® X-100 (both from PanReac AppliChem).

Following blocking, coverslips were incubated overnight at 4°C in a humidified chamber with primary antibodies diluted in the blocking buffer: rabbit anti-pERα (1:500) and mouse monoclonal anti-Caveolin-1 [7C8] (Abcam, ab17052; 1:200). Negative controls were incubated in blocking buffer without primary antibodies. The next day, coverslips were washed in DPBS, then incubated for 1 hour at RT with secondary antibodies diluted in 4% donkey serum (Millipore, #S30) in DPBS. The secondary antibodies used were Alexa Fluor™ 647-conjugated donkey anti-rabbit IgG (1:500) and Alexa Fluor™ 488-conjugated donkey anti-mouse IgG (1:500). Following secondary incubation, coverslips were washed with DPBS and nuclei were counterstained with Hoechst 33342 (Invitrogen, H3570) diluted 1:30,000 in DPBS for 30 minutes at RT. Coverslips were then rinsed with DPBS and Milli-Q water before being mounted onto Superfrost™ Plus adhesion microscope slides (Epredia) using Dako Fluorescence Mounting Medium (Dako, S3023). Slides were stored at 4°C in the dark until imaging.

### Confocal microscopy

Imaging was performed using a Zeiss LSM800 confocal laser scanning microscope equipped with a Plan-Apochromat 63×/1.4 NA oil immersion objective (working distance 0.19 µm). Excitation was provided by three independent laser lines at 405 nm, 488 nm, and 640 nm. The detector gain and laser power were adjusted to avoid saturation or signal clipping. Images were acquired with a pixel resolution of 0.0422553 µm/px at a frame size of 2400 × 2400 pixels. Eight fields of view were imaged per experimental condition, yielding a dataset of 40 images.

### Quantification of spatial distribution of pERα

Image processing and analysis were conducted in FIJI/ImageJ ^79^ to quantify the spatial distribution of pERα across nuclear and cytoplasmic compartments. Nuclear instance image segmentation was carried out using Cellpose 3.0 ^80,81^, employing the pretrained cyto3 model. Raw images were down-sampled to 640 × 640 pixels, and a nucleus radius of 50 pixels was used. Segmentation masks were then upscaled 4× to match the original image dimensions. Cell compartment semantic image segmentation was performed using the LABKIT plugin ^82^ in FIJI on full resolution data. A random forest classifier was trained using manually annotated data from two representative images per experimental group (eight images in total). The classifier categorized pixels into three classes: background, cytoplasm, and nucleus. Image contrast was adjusted to identical levels across the training dataset. Manual annotations were applied to the training images using the Hoechst nuclear staining and the caveolin-1 immunofluorescence signal as proxies for nuclear and cytoplasmic boundaries, respectively. The training process was repeated iteratively until satisfactory segmentation could be verified visually (Fig. S3).

pERα puncta were similarly segmented using a dedicated LABKIT classifier trained on the same annotated image subset (Fig. S3), using the pERα immunofluorescence signal (Alexa Fluor™ 647). Quantification of pERα distribution was performed by computing: (1) the ratio of pERα puncta area to nuclear area in each segmented nucleus, and (2) the ratio of pERα puncta area to total cytoplasmic area per field of view. Boolean AND operations between segmentation masks were applied to define puncta overlapping with nuclear or cytoplasmic ROIs. Nuclei with segmented areas exceeding 10,000 square pixels (17.855 µm²) were retained, while nuclei touching image edges and having small visible areas, were excluded. This area threshold was empirically determined to ensure biologically relevant portions of nuclei were analyzed, given the homogeneous distribution of pERα puncta.

The results from each measurement were compiled in a table and analyzed in Graphpad Prism 10 with a Brown-Forsythe and Welch ANOVA analysis and a Dunnet T3 multiple comparison test to account for potentially different variances across experimental groups.

### Molecular docking and induced-fit docking (IFD)

X-ray crystal structures of the binding domain of ERα (PDB IDs: 1A52 and 5WGD) were retrieved from the RCSB Protein Data Bank <u>(PDB)</u> and were utilized for docking after preparation using the “protein preparation wizard” in Maestro 2021.4 (Schrodinger, LLC, New York, NY, USA). Molecular docking studies were conducted using Glide (Schrodinger, LLC). Compounds were built in the Maestro build panel and optimized to lower energy conformers through LigPrep (Schrodinger, LLC). Hydrogens were added to all atoms of the crystal structure, and the bond orders and formal charges were added to hetero atoms. Chain A was retained, chain B was removed from the asymmetric unit, and side chains not close to the binding pocket or involved in salt bridges were neutralized. Termini were capped with N-acetyl (ACE) and N-methyl amide (NMA) residues. Following preparation, the structure was refined to optimize the hydrogen-bond network using the OPLS_2005 force field. The system was minimized with an RMSD cutoff of 0.30 Å for the convergence of heavy atoms. Prime was utilized to fix possible missing residues/loops or atoms in the protein and to remove co-crystallized water molecules. Water molecules further than 5 Å from the ligand (Hets) were removed. PROPKA was used to check the protonation state of ionizable protein groups (pH = 7.4). Extra precision (XP) docking mode was applied to all compounds on the protein structure grid. Epik was selected as an ionization tool at pH 7.4 ± 1.0. Tautomer generation was also flagged, and the maximum number of conformers generated was set at 32. The ligands of interest were docked with a maximum of five poses for each molecule. The reference ligand, 17β-estradiol, was used as a positive control for interactions with the binding domain of the receptor under analysis. The final assessment of ligand-protein binding was based on the Glide score.

IFD was executed using the IFD module on the sites previously identified, using Maestro 2021.4 (Schrodinger, LLC). The entire receptor molecule was constrained and minimized with an RMSD cutoff of 0.18 Å. Initial Glide docking for each ligand was performed, and side chains were automatically trimmed based on the B-factor, with receptor and ligand van der Waals scaling set to 0.70 and 0.50, respectively. The number of generated poses was set at 10. Prime side-chain prediction and minimization were conducted, refining residues within 5.0 Å of ligand poses and optimizing side chains. This process induced the ligand structure and conformation to fit each pose of the receptor structure. Finally, Glide XP redocking was carried out on structures within 30.0 kcal/mol of the best structure and the top 20 structures overall. The ligand was rigorously docked into the induced-fit receptor structure, yielding an IFD score for each output pose.

### Molecular dynamics (MD) simulations

All MD simulations were performed using GROMACS version 2024.1 ^83^. The Amber99sb force field was employed ^84^ for the estrogen receptor (ERα), while the General Amber Force Field (GAFF) was utilized ^85^ for the ligands: estradiol, S-ketamine, imipramine, fluoxetine, HNK, and PaPE-1. The TIP3P water model was used for solvation ^86^.

### System preparation

Ligand structures (estradiol, S-ketamine, imipramine, fluoxetine, HNK, and PaPE-1) were prepared separately. Partial atomic charges for the ligands were assigned using AM1-BCC as implemented in the Antechamber suite from AmberTools ^87^. Hydrogen atoms were added to the receptor structure using the pdb2gmx utility of GROMACS. The resulting receptor-ligand complex for each of the four ligands formed the starting point for subsequent MD simulations. The following steps were performed independently for each receptor-ligand complex. The receptor-ligand complex was placed in a triclinic simulation box, ensuring a minimum distance of 1.0 nm between the complex and the box edges. The system was then solvated with TIP3P water molecules. To neutralize the system, counter-ions (Na+ or Cl-) were added.

### Energy minimization

The solvated and ionized system underwent energy minimization to remove any steric clashes or unfavorable contacts. This was performed using the steepest descent algorithm for a maximum of 50,000 steps or until the maximum force on any atom was less than 1000.0 kJ mol^-1^ nm^-1^. Long-range electrostatic interactions were treated using the Particle Mesh Ewald (PME) method ^88^ with a cutoff of 1.4 nm. Van der Waals interactions were treated with a cutoff of 1.2 nm, employing a force-switch modifier between 1.0 and 1.2 nm. Periodic boundary conditions (PBC) were applied in all three dimensions.

### Equilibration protocol

A two-stage equilibration protocol was conducted. First, an NVT (canonical ensemble) equilibration was performed for 1 ns (500,000 steps with a 2 fs timestep). During this phase, position restraints were applied to the protein backbone atoms and ligand heavy atoms. The system temperature was maintained at 300 K using the V-rescale thermostat ^89^with a coupling time constant of 0.1 ps. The LINCS algorithm ^90^ was used to constrain bonds involving hydrogen atoms. Second, an NPT (isothermal-isobaric ensemble) equilibration was performed for 1 ns (500,000 steps with a 2 fs timestep). Temperature was maintained at 300 K using the V-rescale thermostat as in the NVT step. Pressure was maintained at 1.0 bar using the Parrinello-Rahman barostat ^91^ with an isotropic coupling type, a time constant of 2.0 ps, and a compressibility of 4.5x10^-5^ bar^-1^. Position restraints on the protein backbone and ligand heavy atoms remained active. All other parameters (constraints, cutoffs, PME, VDW treatment) were identical to the NVT equilibration phase.

### Production MD

Following equilibration, production MD simulations were carried out for 100 ns (50,000,000 steps with a 2 fs timestep) for each receptor-ligand complex under the NPT ensemble. No position restraints were applied during the production run. Temperature (300 K) and pressure (1.0 bar) were maintained using the V-rescale thermostat and Parrinello-Rahman barostat, respectively, with the same parameters as in the NPT equilibration. All other simulation parameters, including PME for long-range electrostatics, Potential-shift-Verlet for VDW interactions (cutoffs at 1.0 nm), and LINCS for H-bond constraints, were consistent with the NPT equilibration phase. Trajectory coordinates were saved every 2 ps (1000 steps).

### Trajectory analysis

The production MD trajectories were analyzed using GROMACS utilities. Prior to analysis, trajectories were corrected for PBC, centered, and fitted to the protein backbone to remove overall rotational and translational movements.

The stability of the systems was assessed by calculating the RMSD of the protein backbone atoms and ligand heavy atoms (fitted to the protein backbone) relative to their initial equilibrated structures. The root mean square fluctuation (RMSF) per residue was calculated for the protein Cα atoms to identify flexible regions. The radius of gyration (RoG) of the protein was computed to evaluate its compactness. The solvent-accessible surface area (SASA) of the protein-ligand complex was also calculated.

### Binding free energy calculations

The binding free energies for each receptor-ligand complex were estimated using the Molecular Mechanics/Poisson-Boltzmann Surface Area (MM/PBSA) and Molecular Mechanics/Generalized Born Surface Area (MM/GBSA) methods as implemented in the gmx_MMPBSA tool (version 1.6.4) ^92^. Calculations were performed on 161 frames extracted from the last 80 ns of the production MD trajectory, with an interval of 250 frames (corresponding to snapshots taken every 0.5 ns from 20 ns to 100 ns). For MM/GBSA calculations, the GB model igb = 5 was used with an internal dielectric constant of 1.0 and an external dielectric constant of 78.5.

For MM/PBSA calculations, the Poisson-Boltzmann equation was solved using ipb = 2, with an internal dielectric constant (indi) of 1.0 and an external dielectric constant (exdi) of 80.0. Atomic radii were set using radiopt = 1 (Bondi radii). Ionic strength was set to 0.0 M for both calculations.

### Proximity ligation assay (PLA) for ERα–antidepressant interaction

MCF-7 and HEK293T cells were used. HEK293T cells were seeded on 13 mm poly-L-lysine-coated glass coverslips, no coating was performed for MCF-7 cells. A group of HEK293T cells were transfected with a plasmid encoding ERα (pEGFP-C1-ER alpha, Addgene, Plasmid #28230) using Lipofectamine 2000 (Thermo Fisher Scientific). Non-transfected cells served as a negative control. After 24 hours, the media were replaced and cells were treated with 10 µM biotinylated fluoxetine and HNK (using EZ-Link NHS-PEG4 Biotinylation Kit, Thermo Scientific, #21455, following manufactureŕs protocol) for 10 minutes at 37°C, then fixed with 4% PFA for 15 minutes. Cells were washed with PBS and stored at 4°C until further processing.

PLA was performed according to the manufacturer’s instructions with minor modifications using Duolink® In Situ Red Starter Kit Mouse/Rabbit (DUO92101-1KT, Sigma-Aldrich). After removing PBS, coverslips were incubated with Duolink® Blocking Solution for 1 hour at 37°C. Streptavidin-HRP (Thermo Scientific, #21126) (1:5000 in Antibody Diluent) was applied for 45 minutes at room temperature, followed by two washes with Wash Buffer A. Coverslips were then incubated overnight at 4°C with mouse anti- ERα (Enzo Life Sciences, ADI-SRA-1010) (1:1000) and rabbit anti-HRP (Rockland, #200-4138) (1:1000) in Antibody Diluent. After two additional washes, PLUS and MINUS PLA probes (1:5 dilution) were added and incubated for 1 hour at 37°C. Ligation was performed with freshly prepared ligation buffer and ligase (1:40 dilution), followed by a 30-minute incubation at 37°C. Slides were washed and subjected to amplification using polymerase (1:80 dilution) in amplification buffer for 100 minutes at 37°C in the dark. Coverslips were washed twice in Wash Buffer B and once in 0.01× Wash Buffer B, then mounted with Duolink® In Situ Mounting Medium containing DAPI. Z-stack images (≥15 slices, 1.5 µm) were acquired using a 63× oil objective with 3× zoom on a Zeiss LSM800 confocal microscope. Blue (ex358/em465 nm) and red (ex594/em624 nm) channels were used to detect DAPI and PLA signals, respectively.

### Statistical analysis

Statistical analyses were conducted using GraphPad Prism (10.5.0). For comparisons involving more than two groups, one-way ANOVA was employed and paired or unpaired student t-tests were performed to compare two groups. To see the interaction effects, 2-way ANOVA was used. Tukey’s and Dunn’s multiple comparison tests were applied. In Western blotting, technical replicates of the three biological replicates were pooled. Whenever necessary, data were log-transformed to meet the assumption of homoscedasticity. For the colocalization assay, a Brown-Forsythe and Welch ANOVA was used to account for variance heterogeneity, followed by Dunnett’s T3 multiple comparisons test to assess group-wise differences. Error bars on the graphs represent mean ± SEM.

## Data availability

Data will be available upon request to the corresponding authors.

## Author Contributions

**SA:** Conceptualization, Investigation, Methodology, Formal analysis, Data curation, Visualization, Writing - Original draft, Writing - Review & editing, Funding acquisition; **MR:** Investigation, Formal analysis, Writing - original draft; **DS:** Formal analysis, Visualization; **CC:** Investigation; **RR:** Investigation, **CBV:** Resources, Writing - Review & editing; **HKM:** Resources, Supervision, Writing - Review & editing; **JS:** Investigation; **MNS:** Resources, Writing - Review & editing; **US:** Resources, Writing - Review & editing; **SS:** Conceptualization, Visualization, Resources, Supervision, Writing - Review & editing, **AML:** Supervision, Writing - Review & editing, **GW:** Conceptualization, Resources, Supervision, Writing - Review & editing, Funding acquisition; **SJ:** Resources, Supervision, Conceptualization, Writing - Review & editing; **CB:** Conceptualization, Investigation, Methodology, Supervision, Writing - Review & editing.

## Acknowledgment

The authors thank Plinio Casarotto for his valuable comments on the experimental design; Birgit Schiøtt for granting S.A. access to the Schrödinger Suite, and <u>Schrödinger</u>, Inc. for providing a trial license for complementary analysis; Nicole Silva, Judit Prat Duran, Fernanda Crunfli, Erik Kaadt, Amanda Dyrholm Strange, Marie Vadstrup Pedersen, and Malene Overby for stimulating discussions; Daniela Grimm for providing various batches of the cell line used in the study. The authors acknowledge the Bioimaging Core Facility, Health, Aarhus University, Denmark, for the use of equipment and support.

