## Supplementary material for "Antidepressants interact with sex steroid receptors and their intracellular signaling components": Supplementary materials_20260222.docx

^*^Correspondence to:

Fig. S1

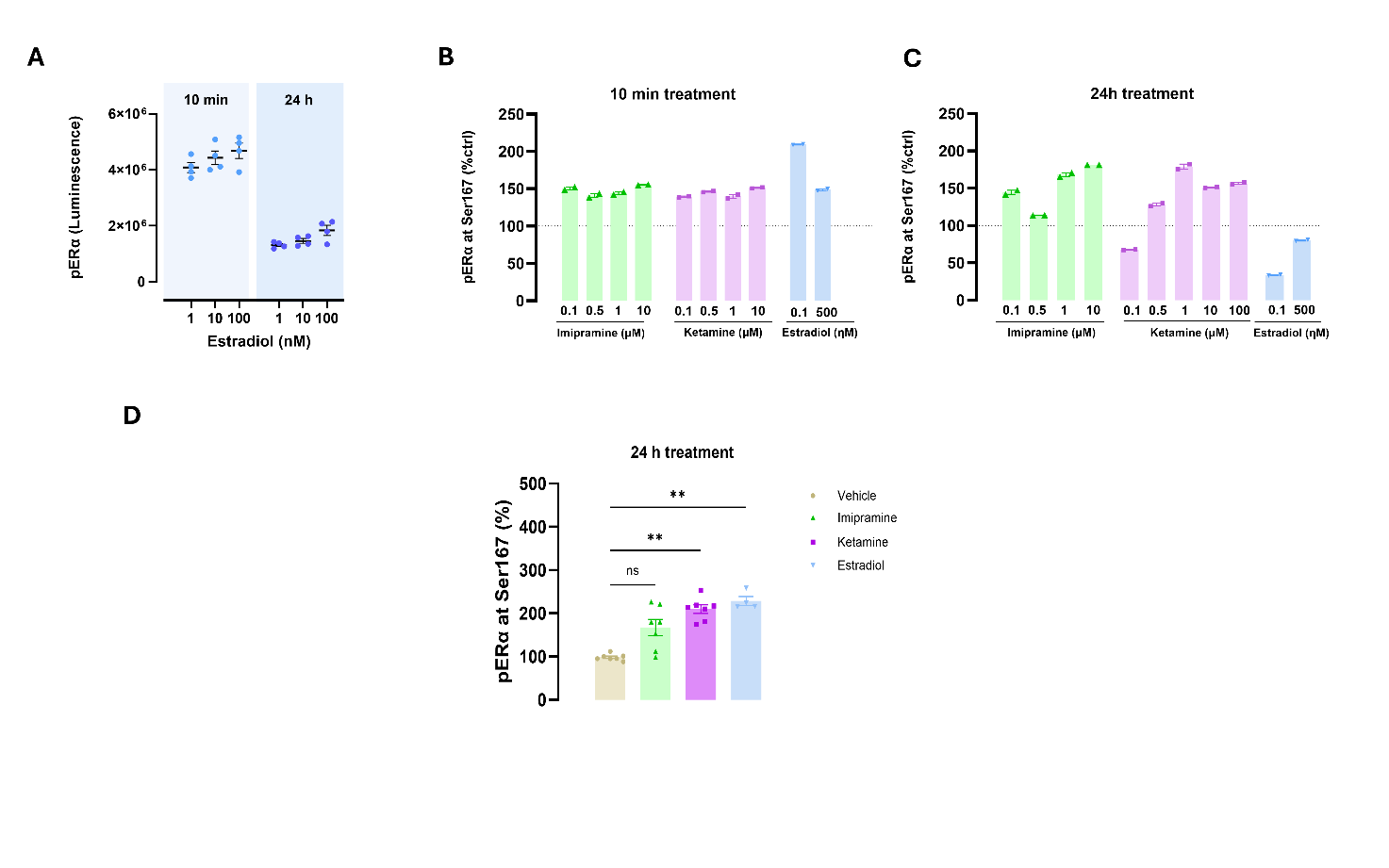

Fig. S1 |

(A) Dose-dependent induction of ERα Ser167 phosphorylation by estradiol (E2) following acute (10 min) and chronic (24 h) treatment in MCF-7 cells. (B and C) Dose–response analysis of antidepressants identifies imipramine and S-ketamine as effective inducers of ERα Ser167 phosphorylation; both compounds were used at 10 µM in subsequent experiments. (D) Chronic (24 h) treatment with E2, imipramine and S-ketamine increases ERα Ser167 phosphorylation (p < 0.01 for E2 and S-ketamine and p = 0.0731 for imipramine).

Fig. S2

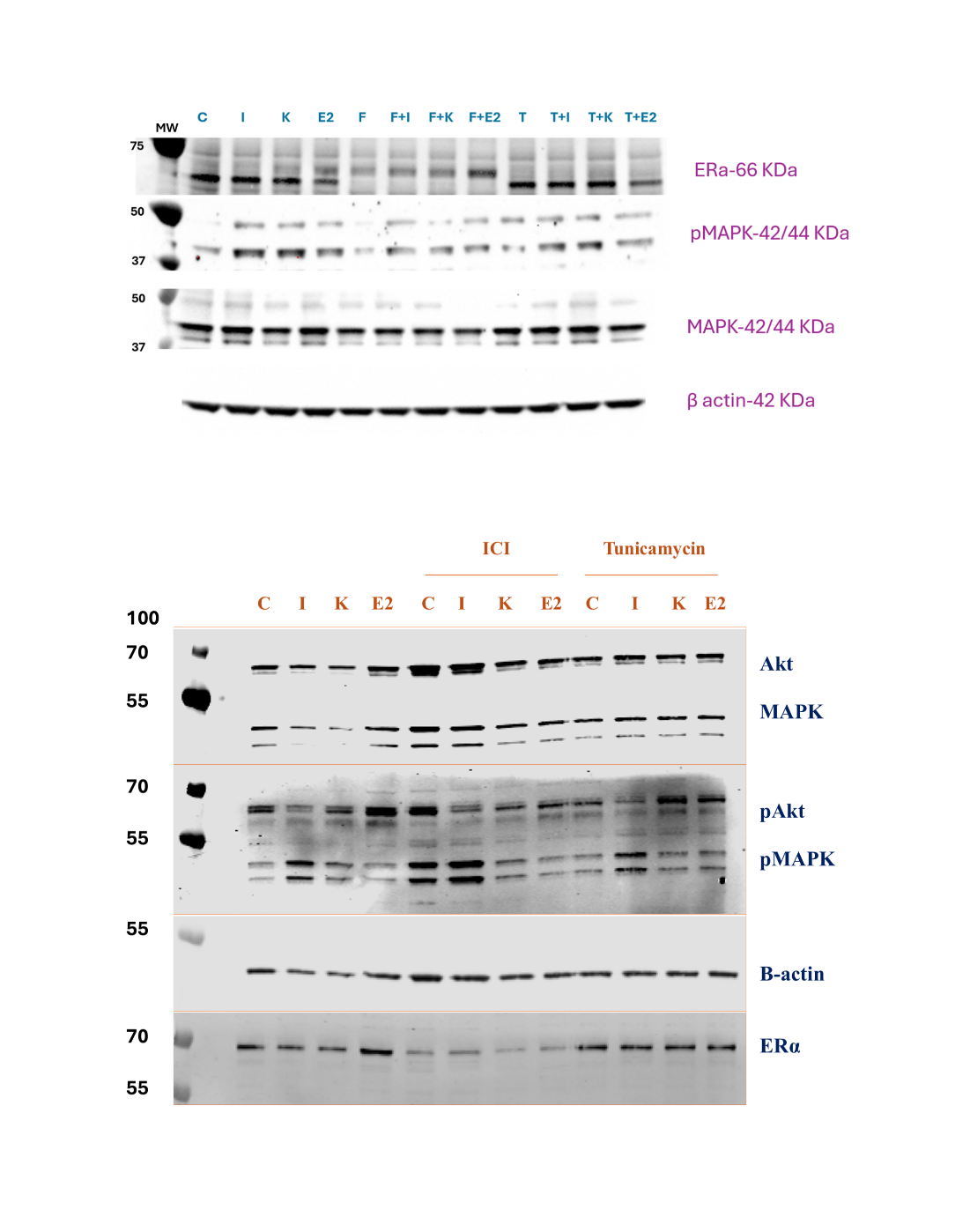

Fig. S2 |

Representative Western blots of whole-cell lysates from MCF-7 cells treated with vehicle (C), 17β-estradiol (E2), imipramine (I), or S-ketamine (K), probed with the indicated antibodies.

Table S1 | Binding profiles of different antidepressants with the ligand binding domain of ERα.

| Ligands | Docking score | Glide  ligand  efficiency | Glide  ligand  efficiency SA | Glide  ligand  efficiency ln | XP GScore | Glide gscore | Glide energy | Glide  einternal | Glide emodel | XP HBond |
| --- | --- | --- | --- | --- | --- | --- | --- | --- | --- | --- |
| Duloxetine | -11.772 | -0.561 | -1.547 | -2.911 | -11.772 | -11.772 | -42.434 | 3.188 | -58.507 | -0.871 |
| Paroxetine | -11.075 | -0.461 | -1.331 | -2.651 | -11.078 | -11.078 | -34.573 | 9.393 | -49.183 | 0 |
| Fluoxetine | -10.693 | -0.486 | -1.362 | -2.614 | -10.693 | -10.693 | -30.818 | 3.958 | -50.444 | -0.9 |
| Milnacipran | -9.743 | -0.541 | -1.419 | -2.504 | -9.746 | -9.746 | -25.879 | 6.951 | -37.929 | -0.557 |
| Modafinil | -9.368 | -0.493 | -1.316 | -2.375 | -9.368 | -9.368 | -41.179 | 2.648 | -59.938 | -0.972 |
| Imipramine | -9.229 | -0.439 | -1.212 | -2.282 | -9.229 | -9.229 | -32.904 | 4.034 | -47.049 | 0 |
| HNK | -8.949 | -0.559 | -1.409 | -2.372 | -9.184 | -9.184 | -33.339 | 9.041 | -41.925 | -1.141 |
| Venlafaxine | -8.628 | -0.431 | -1.171 | -2.159 | -8.63 | -8.63 | -27.438 | 4.865 | -35.54 | -0.304 |
| Ketamine | -8.203 | -0.513 | -1.292 | -2.174 | -8.448 | -8.448 | -29.207 | 6.677 | -37.178 | 0 |
| Cannabidiol | -7.827 | -0.34 | -0.968 | -1.893 | -7.827 | -7.827 | -12.834 | 3.712 | 17.467 | 0 |
| Citalopram | -7.285 | -0.304 | -0.876 | -1.744 | -7.29 | -7.29 | -35.641 | 8.405 | -25.421 | -0.35 |
| Nortriptyline | -6.828 | -0.341 | -0.927 | -1.709 | -6.828 | -6.828 | -32.584 | 2.708 | -41.222 | 0 |
| Psilocin | -6.779 | -0.452 | -1.115 | -1.828 | -6.785 | -6.785 | -29.053 | 1.572 | -40.483 | -0.7 |
| Maprotiline | -6.662 | -0.317 | -0.875 | -1.647 | -6.662 | -6.662 | -27.891 | 3.392 | -23.525 | 0 |
| Bupropion | -6.367 | -0.398 | -1.003 | -1.688 | -6.702 | -6.702 | -24.703 | 9.506 | -25.222 | 0 |
| Mianserin | -6.305 | -0.315 | -0.856 | -1.578 | -6.644 | -6.644 | -30.676 | 0 | -7.307 | 0 |
| Mirtazapine | -6.261 | -0.313 | -0.85 | -1.567 | -6.455 | -6.455 | -32.683 | 0 | -0.838 | 0 |
| Vortioxetine | -5.77 | -0.275 | -0.758 | -1.427 | -5.787 | -5.787 | -34.42 | 4.56 | -49.263 | 0 |
| Sertraline | -5.737 | -0.287 | -0.779 | -1.436 | -5.769 | -5.769 | -23.79 | 4.704 | -33.638 | 0 |
| Fluvoxamine | -4.747 | -0.216 | -0.605 | -1.16 | -4.757 | -4.757 | -39.281 | 7.799 | -50.81 | 0 |
| Psilocybin | -2.231 | -0.117 | -0.313 | -0.566 | -2.231 | -2.231 | -29.916 | 4.199 | -40.715 | 0 |

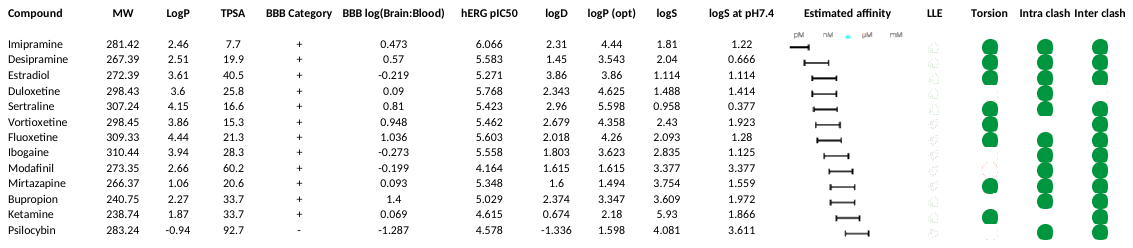
 Table S2 | Estimated binding affinity of compounds for ERα and predicted physicochemical properties relevant to CNS drug delivery.

**MW**: molecular weight; **logP**: Logarithm of n-octanol-water partition coefficient, describing the relationship between lipophilicity and hydrophilicity of a neutral compound; **TPSA**: Topological Polar Surface Area; **BBB log(Brain:Blood)**: Logarithm of blood-brain partition coefficient of a compound; **hERG pIC50**: Prediction of a drug's pIC50 values in blocking the human Ether-à-go-go-Related Gene (hERG) potassium channel, indicating potential for cardiotoxicity; **logD:** Logarithm of n-octanol-water partition coefficient at the physiological pH7.4, describing the relationship between lipophilicity and hydrophilicity of an ionized compound; **logS**: Logarithm of intrinsic aqueous solubility in μM for neutral compound; **logS at pH 7.4**: Logarithm of intrinsic aqueous solubility at physiological pH 7.4 in μM for ionizable compounds; **LLE**: Ligand-lipophilicity efficiency predicted by HYDE based on desolvation and interactions.

**A**
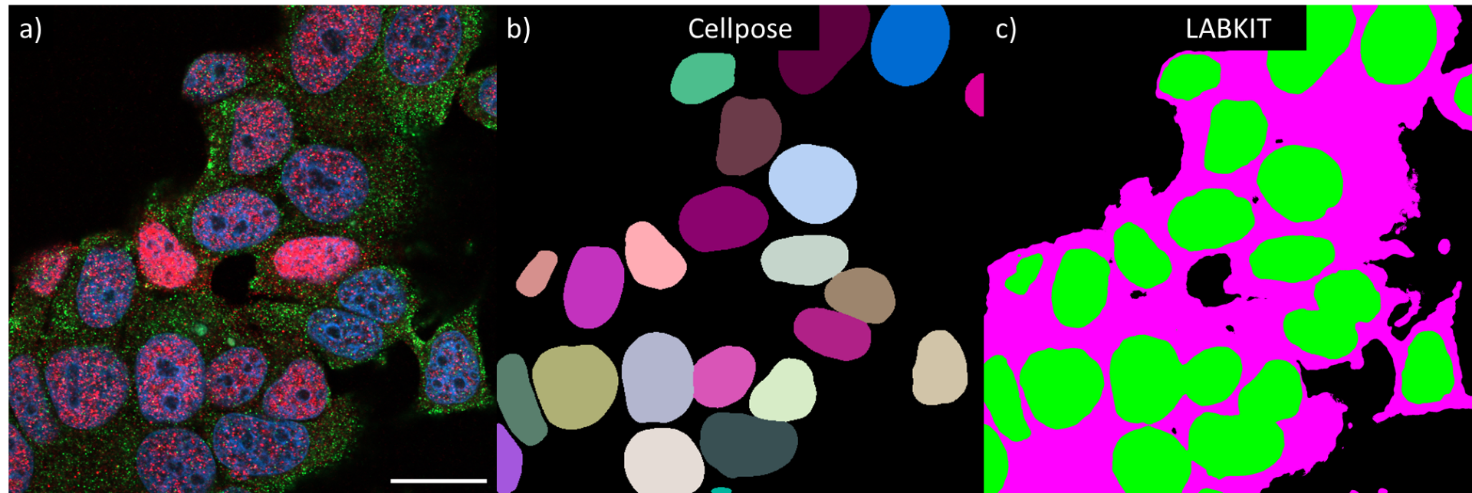

**B**
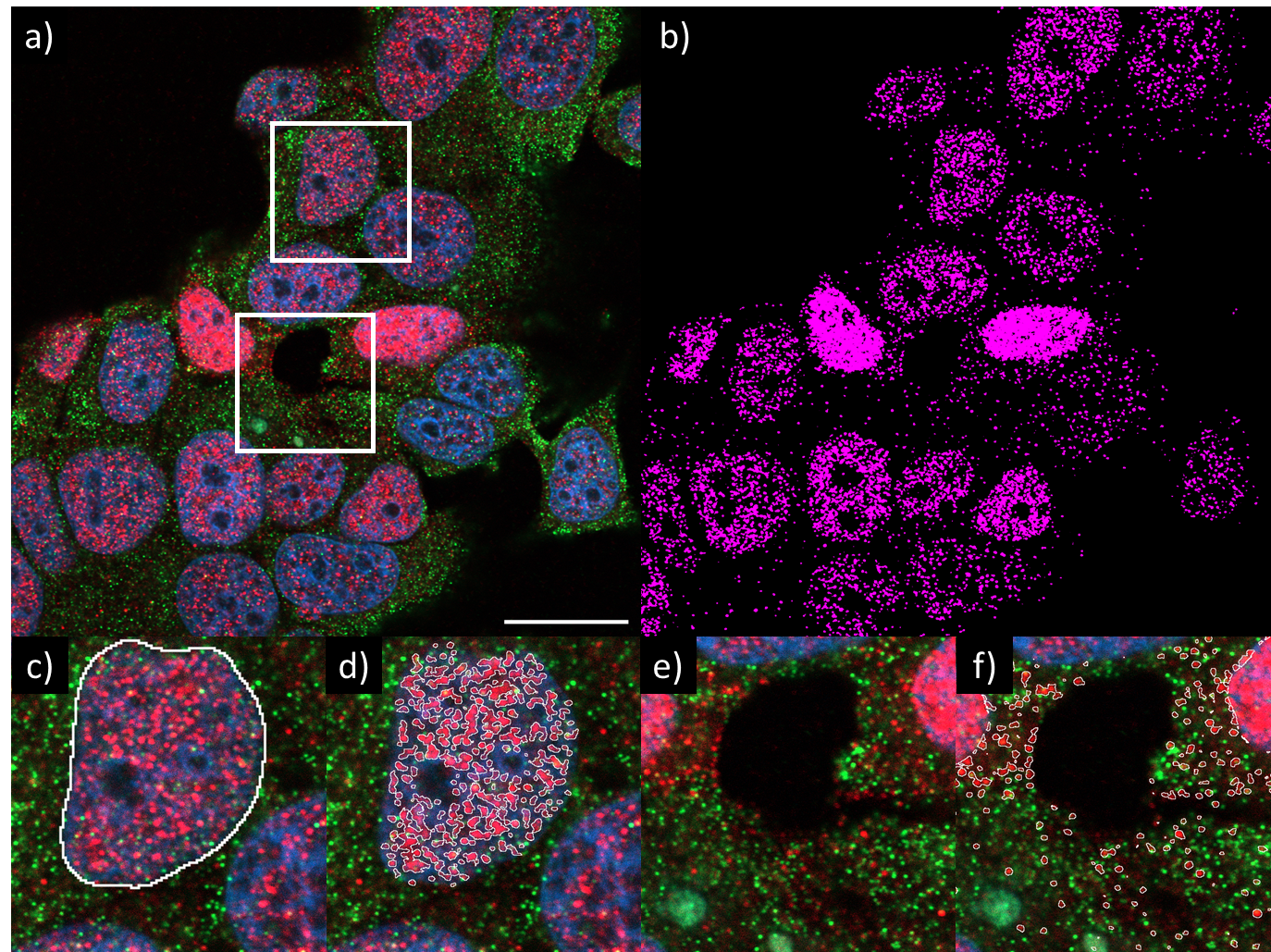

**Fig S3 | Segmentation of cellular compartments and pERα puncta.**

(A) Segmentation of cellular compartments. a) Raw data from imipramine-treated group. b) Instance segmentation of cell nuclei with Cellpose, every nucleus has a unique numerical identifier. c) LABKIT Semantic segmentation of cytoplasm (magenta) from background (black) and nuclei (green). Scale bar = 20 µm.

(B) Segmentation of pERα puncta. a) Raw data from Imipramine-treated group. b) LABKIT semantic segmentation of cell puncta. c-d) Segmentation of pERα puncta within each nucleus using Cellpose instance segmentation images. Puncta are quantified on a per nucleus basis. e-f) Segmentation of PLA puncta in cytoplasm using the LABKIT segmentation map. Scale bar= 20µm.
